# A tRNA processing enzyme is a central regulator of the mitochondrial unfolded protein response

**DOI:** 10.1101/2021.06.22.449331

**Authors:** James P. Held, Benjamin R. Saunders, Claudia V. Pereira, Maulik R. Patel

## Abstract

The mitochondrial unfolded protein response (UPR^mt^) has emerged as a predominant mechanism that preserves mitochondrial function. Consequently, multiple pathways likely exist to modulate UPR^mt^. We unexpectedly discovered that the tRNA processing enzyme, homolog of ELAC2 (HOE-1), is central to UPR^mt^ regulation in *Caenorhabditis elegans*. We find that nuclear HOE-1 is necessary and sufficient to robustly activate UPR^mt^. We show that HOE-1 acts via transcription factors ATFS-1 and DVE-1 that are crucial for UPR^mt^. Mechanistically, we show that HOE-1 likely mediates its effects via tRNAs, as blocking tRNA export prevents HOE-1-induced UPR^mt^. Interestingly, we find that HOE-1 does not act via the integrated stress response, which can be activated by uncharged tRNAs, pointing towards its reliance on a new mechanism. Finally, we show that the subcellular localization of HOE-1 is responsive to mitochondrial stress and is subject to negative regulation via ATFS-1. Together, we have discovered a novel RNA-based cellular pathway that modulates UPR^mt^.

---

Mitochondria are central to a myriad of cellular processes including energy production, cellular signaling, biogenesis of small molecules, and regulation of cell death via apoptosis ^1^. Given the importance of mitochondria, it is no surprise that mitochondrial dysfunction can lead to metabolic and neurological disorders, cardiovascular disease, and cancers ^2^. To maintain proper mitochondrial function cellular mechanisms have evolved that respond to, and mitigate, mitochondrial stress ^3–11^.

One of the predominant mitochondrial stress response mechanisms is the mitochondrial unfolded protein response (UPR^mt^). Though first discovered in mammals ^12^, UPR^mt^ has been best characterized in *Caenorhabditis elegans* ^9^. UPR^mt^ is primarily characterized by transcriptional upregulation of genes whose products respond to and ameliorate mitochondrial stress ^13,14^.

In *C. elegans*, activation of UPR^mt^ is dependent upon the transcription factor ATFS-1 which primarily localizes to mitochondria, but under mitochondrial-stress conditions is trafficked to the nucleus where it drives the expression of mitochondrial stress response genes ^14–16^. However, it has become increasingly apparent that UPR^mt^ is under multiple levels of control: Mitochondrial stress in neurons can activate intestinal UPR^mt^ non-cell-autonomously via retromer-dependent Wnt signaling ^17–19^; overexpression of two conserved histone demethylases are independently sufficient to activate UPR^mt^ ^20^; and ATFS-1 is post-translationally modified to facilitate its stability and subsequent UPR^mt^ activation ^21^. Given its importance, there are likely yet-to-be identified pathways regulating UPR^mt^.

We serendipitously discovered that the 3’-tRNA zinc phosphodiesterase, homolog of ELAC2 (HOE-1) is a key regulator of UPR^mt^ in *C. elegans*. ELAC2 is an essential endonuclease that cleaves 3’-trailer sequences from nascent tRNAs—a necessary step of tRNA maturation—in both nuclei and mitochondria ^22–29^. ELAC2 has also been reported to cleave other structured RNAs yielding tRNA fragments, small nucleolar RNAs (snoRNAs) and micro RNAs (miRNAs) ^29–32^. In humans, mutations in ELAC2 are associated with hypertrophic cardiomyopathy ^33–35^ and prostate cancer ^36–38^ while in *C. elegans*, loss of HOE-1 has been shown to compromise fertility ^39^.

We find that nuclear HOE-1 is required for activation of UPR^mt^. Remarkably, compromising nuclear export of HOE-1 is sufficient to specifically and robustly activate UPR^mt^, even in the absence of mitochondrial stress. Blocking tRNA export from the nucleus suppresses this HOE-1-dependent UPR^mt^ induction, suggesting that HOE-1 generates RNA species required in the cytosol to trigger UPR^mt^. Finally, we show that HOE-1 nuclear levels are dynamically regulated under conditions of mitochondrial stress, supporting a physiological role for HOE-1 in mitochondrial stress response. Taken together, our results provide a novel mechanism by which UPR^mt^ is regulated as well as provide critical insight into the biological role of HOE-1.

## Results

### *hoe-1* is required for maximal UPR^mt^ activation

We discovered that RNAi against *hoe-1*, a gene encoding a 3’-tRNA phosphodiesterase, attenuates *hsp-6p::GFP* induction—a fluorescence based transcriptional reporter of UPR^mt^ activation ^13^. Knockdown of *hoe-1* by RNAi is sufficient to attenuate UPR^mt^ reporter activation induced by a loss-of-function mutation in the mitochondrial electron transport chain (ETC) complex I subunit NUO-6 (*nuo-6(qm200)*) **(Fig. 1a, b)**.

**Fig. 1:**
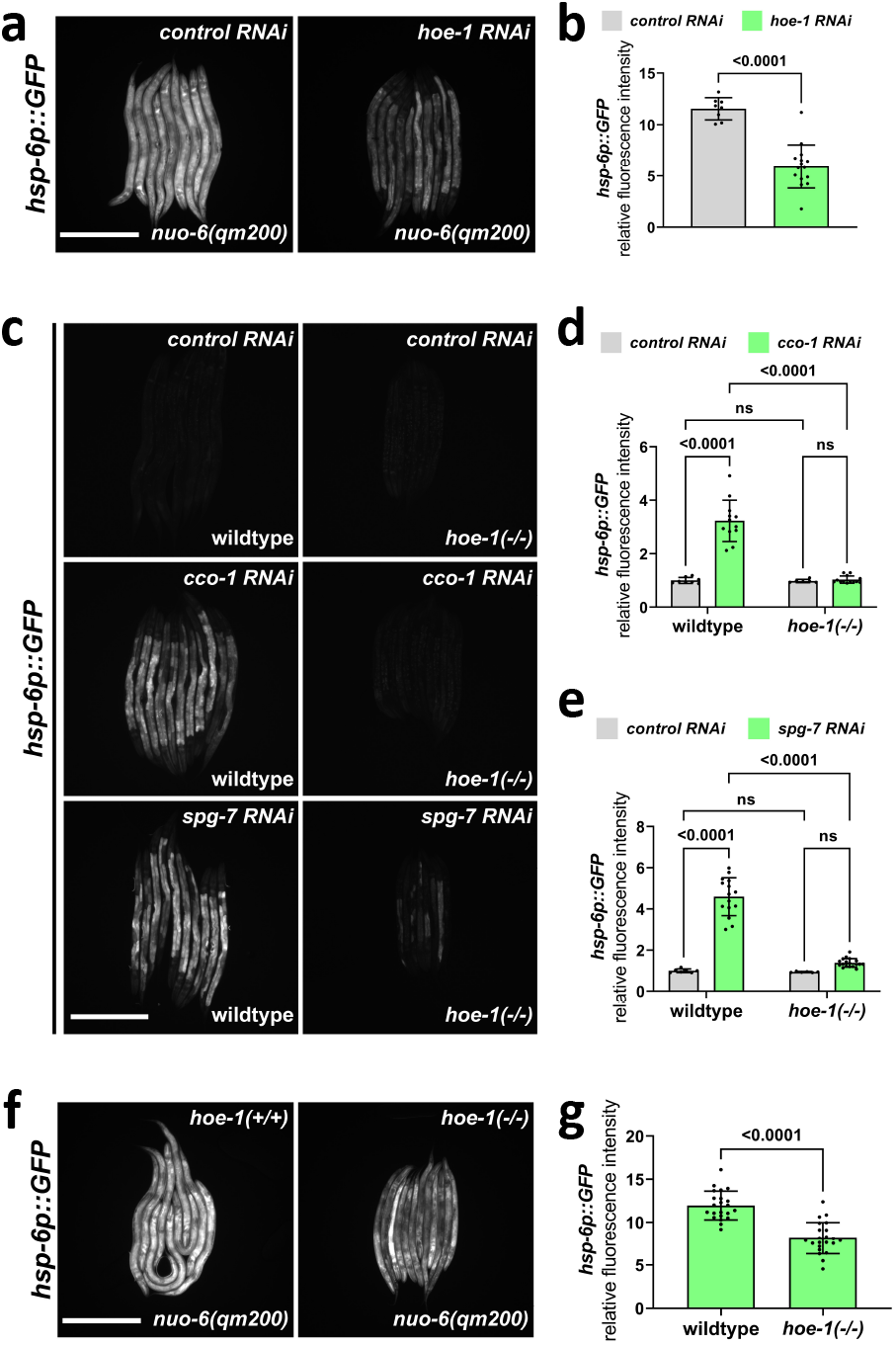
*hoe-1* is required for maximal UPR^mt^ activation. **a**, Fluorescence images of UPR^mt^ reporter (*hsp-6p::GFP)* activation in L4 *nuo-6(qm200)* animals on control and *hoe-1* RNAi. Scale bar 200μm. **b**, Fluorescence intensity quantification of *hsp-6p::GFP* in individual L4 *nuo-6(qm200)* animals on control and *hoe-1* RNAi normalized to *hsp-6p::GFP* in a wildtype background (n=8 and 15 respectively, mean and SD shown, two-tailed unpaired t-test). **c**, Fluorescence images of UPR^mt^ reporter (*hsp-6p::GFP*) activation in L3/L4 wildtype and *hoe-1* null (*hoe-1(−/−)*) animals on control, *cco-1*, and *spg-7* RNAi. Scale bar 200μm. **d**, Fluorescence intensity quantification of *hsp-6p::GFP* in individual L3/L4 wildtype and *hoe-1(−/−)* animals on control and *cco-1 RNAi* (n=8,12,6 and 13 respectively, mean and SD shown, ordinary two-way ANOVA with Tukey’s multiple comparisons test). **e**, Fluorescence intensity quantification of *hsp-6p::GFP* in individual L3/L4 wildtype and *hoe-1(−/−)* animals on control and *spg-7 RNAi* (n=7,15,6 and 18 respectively, mean and SD shown, ordinary two-way ANOVA with Tukey’s multiple comparisons test). **f**, Fluorescence images of UPR^mt^ reporter (*hsp-6p::GFP*) activation in L3/L4 *nuo-6(qm200)* animals with wildtype *hoe-1* (*hoe-1(+/+)*) or *hoe-1* null (*hoe-1(−/−)*). Scale bar 200μm. **g**, Fluorescence intensity quantification of *hsp-6p::GFP* in individual L3/L4 *nuo-6(qm200)* animals with *hoe-1(+/+)* or *hoe-1(−/−)* normalized to *hsp-6p::GFP* in a wildtype background (n=22 for each condition, mean and SD shown, two-tailed unpaired t-test).

To further interrogate the potential role of *hoe-1* in UPR^mt^ regulation we used CRISPR/*Cas9* to generate a *hoe-1* null mutant by deleting the open reading frame of *hoe-1*. The *hoe-1* null mutants do not develop past late larval stage 3, thus the allele is maintained over a balancer chromosome, *tmC25* ^40^. UPR^mt^ induced by the knockdown of both the mitochondrial protease, *spg-7*, and ETC complex IV subunit, *cco-1*, is robustly attenuated in *hoe-1* null animals **(Fig. 1c, d, e)**. Furthermore, UPR^mt^ induced by *nuo-6(qm200)* is attenuated in *hoe-1* null animals similarly to what is seen in *nuo-6(qm200)* animals on *hoe-1* RNAi **(Fig. 1f, g)**. Taken together, these findings suggest that HOE-1 is generally required for maximal UPR^mt^ activation.

### Nuclear HOE-1 is required for maximal UPR^mt^ activation

To better understand the role of HOE-1 in UPR^mt^ regulation we sought to identify where HOE-1 functions in the cell. HOE-1 is predicted to localize to both nuclei and mitochondria and this dual-localization has been shown for HOE-1 orthologs in *Drosophila*, mice, and human cell lines ^26,27,29,41^. To determine the subcellular localization of HOE-1 in *C. elegans* we tagged HOE-1 with GFP at its endogenous locus (HOE-1::GFP). We found that HOE-1 localizes to both mitochondria and nuclei **(Fig. 2a)**.

**Fig. 2:**
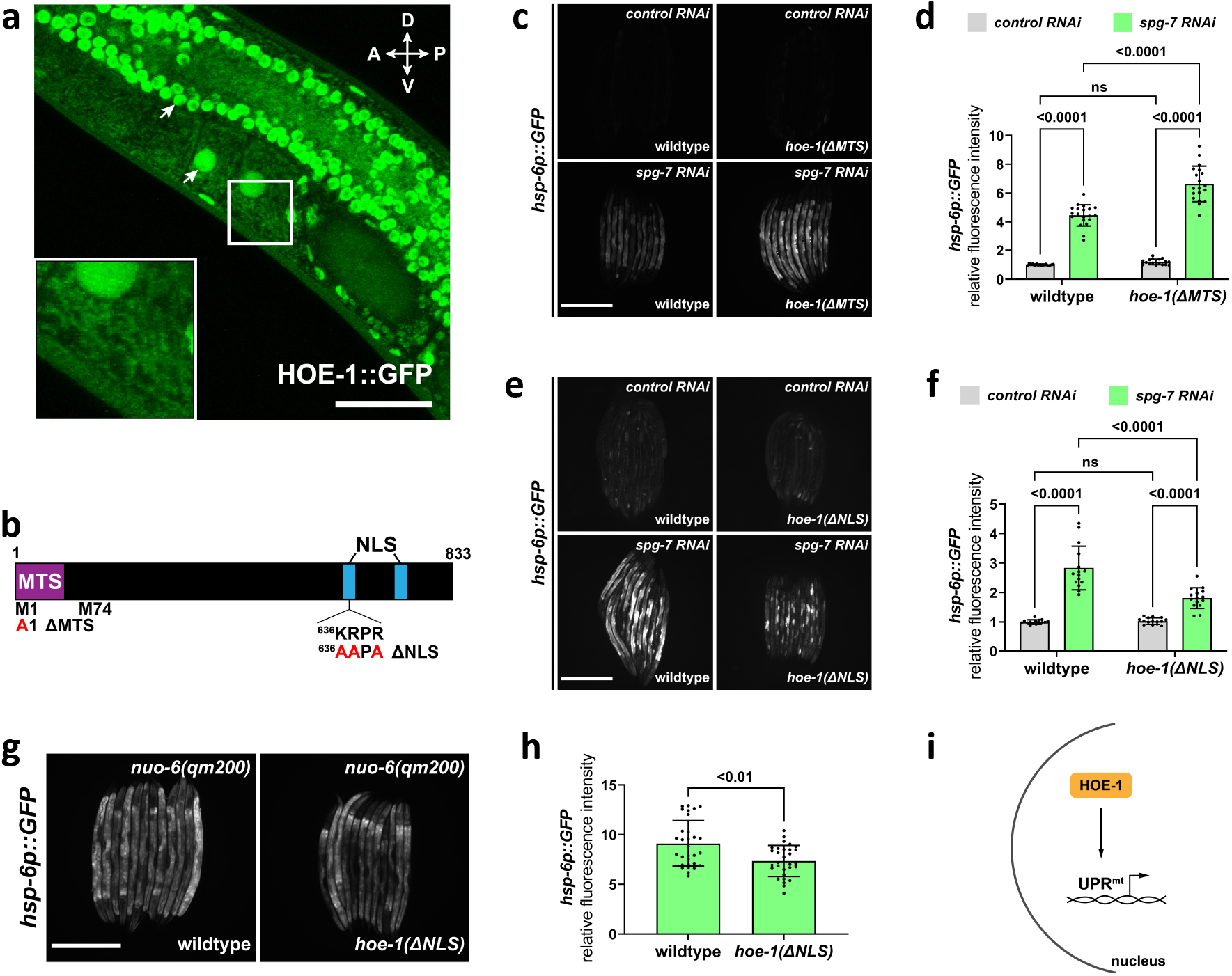
Nuclear HOE-1 is required for maximal UPR^mt^ activation. **a**, Partial z-stack maximum intensity projection fluorescence image of HOE-1::GFP in the adult germline. Orientation compass: anterior (A), posterior (P), dorsal (D), ventral (V). Arrows indicate nuclei. Area in the white square magnified in inset to show mitochondria. Scale bar 25μm. **b**, Schematic of HOE-1 protein showing the mitochondrial targeting sequence (MTS) and nuclear localization signals (NLS). ΔMTS allele created by replacing START codon with an alanine (M1A). Transcription begins at M74 for nuclear localized HOE-1. ΔNLS allele created by compromising the most N-terminal NLS (^636^KRPR > AAPA). **c**, Fluorescence images of UPR^mt^ reporter (*hsp-6p::GFP*) in L4 wildtype and *hoe-*1(ΔMTS) animals on control and *spg-7 RNAi*. Scale bar 200μm. **d**, Fluorescence intensity quantification of *hsp-6p::GFP* in individual L4 wildtype and *hoe-1*(ΔMTS) animals on control and *spg-7 RNAi* (n=15,20,17, and 19 respectively, mean and SD shown, ordinary two-way ANOVA with Tukey’s multiple comparisons test). **e**, Fluorescence images of UPR^mt^ reporter (*hsp-6p::GFP*) in L4 wildtype and *hoe-*1(ΔNLS) animals on control and *spg-7 RNAi*. Scale bar 200μm. **f**, Fluorescence intensity quantification of *hsp-6p::GFP* in individual L4 wildtype and *hoe-1*(ΔNLS) animals on control and *spg-7 RNAi* (n=15 for each condition, mean and SD shown, ordinary two-way ANOVA with Tukey’s multiple comparisons test). **g**, Fluorescence images of UPR^mt^ reporter in L4 *nuo-6(qm200)* animals in wildtype and *hoe-1*(ΔNLS) backgrounds. Scale bar 200μm. **h**, Fluorescence intensity of *hsp-6p::GFP* in individual L4 *nuo-6(qm200)* animals in wildtype and *hoe-1*(ΔNLS) backgrounds (n=30 for each condition, mean and SD shown, two-tailed unpaired t-test). **i**, Schematic depicting the requirement of HOE-1 in the nucleus for complete UPR^mt^ activation.

Given the dual-localization of HOE-1, we questioned whether it is mitochondrial or nuclear HOE-1 that is required for UPR^mt^ activation. To address this question, we created mitochondrial and nuclear compartment-specific knockouts of HOE-1 **(Fig. 2b)**. *hoe-1* contains two functional start codons. Transcription beginning from the first start codon (encoding methionine 1 (M1)) produces mRNA that encodes a mitochondrial targeting sequence (MTS). Transcription beginning from the second start codon (encoding methionine 74 (M74)), which is 3’ to the MTS, produces a nuclear specific transcript. This feature is conserved in human ELAC2 and it has been shown that mutating M1 to an alanine produces a mitochondrial-specific knockout ^27^. Thus, we used the same approach to create a mitochondrial-specific knockout of *C. elegans* HOE-1 (*hoe-1*(ΔMTS)). This mutation is sufficient to ablate mitochondrial targeting without impacting nuclear localization **(Ext. Data Fig. 1)**. HOE-1 is predicted to contain two nuclear localization signals (NLS). To ablate nuclear localization we mutated the positively charged residues of the most N-terminal NLS to alanine (*hoe-1*(ΔNLS)). These mutations are sufficient to ablate HOE-1 nuclear localization **(Ext Data Fig. 2)**.

UPR^mt^ reporter activation by *spg-7* or *cco-1* RNAi is not attenuated in *hoe-1*(ΔMTS) animals **(Fig. 2c, d and Ext. Data Fig. 3a, b)**. In fact, UPR^mt^ reporter activation is slightly elevated in *hoe-1*(ΔMTS) animals relative to wildtype. In contrast, loss of nuclear HOE-1 robustly attenuates UPR^mt^ activation by *spg-7* RNAi like that seen in *hoe-1* null animals **(Fig. 2e, f)**. Furthermore, loss of nuclear HOE-1 attenuates UPR^mt^ activation induced by *nuo-6(qm200)* **(Fig. 2g, h)**. Together these data suggest that HOE-1 is required in the nucleus to facilitate UPR^mt^ activation **(Fig. 2i)**.

### Compromising HOE-1 nuclear export is sufficient to activate UPR^mt^

Like many nuclear localized proteins ^42^, HOE-1 has both nuclear localization signals and a nuclear export signal (NES). Given that loss of nuclear HOE-1 results in UPR^mt^ attenuation we questioned if compromising HOE-1 nuclear export, by ablating the NES of HOE-1, is sufficient to activate UPR^mt^. We created a HOE-1 NES knockout mutant (*hoe-1*(ΔNES)) by replacing the strong hydrophobic resides of the predicted NES with alanines **(Ext. Data Fig. 4a)**. *hoe-1*(ΔNES) animals are superficially wildtype in their development but are sterile. Thus, the allele is balanced with *tmC25*.

Strikingly, the UPR^mt^ reporter *hsp-6p::GFP* is robustly activated, predominantly in the intestine, in *hoe-1*(ΔNES) animals. Induction of *hsp-6p::GFP* by *hoe-1*(ΔNES) begins in larval stage 4 (L4) animals **(Ext. Data Fig. 4b, c)** and is robustly activated by day 2 of adulthood **(Fig. 3a, b)**. Induction of *hsp-6p::GFP* by *hoe-1*(ΔNES) in day 2 adults is more robust than induction by the mitochondrial stressor *nuo-6(qm200)* and as robust as constitutive UPR^mt^ activation by an *atfs-1* gain-of-function mutation (*atfs-1(et15)*) **(Fig. 3a, b)**. *hoe-1*(ΔNES) also mildly induces the less sensitive UPR^mt^ reporter *hsp-60p::GFP* **(Ext. Data Fig. 4d, e)**.

**Fig. 3:**
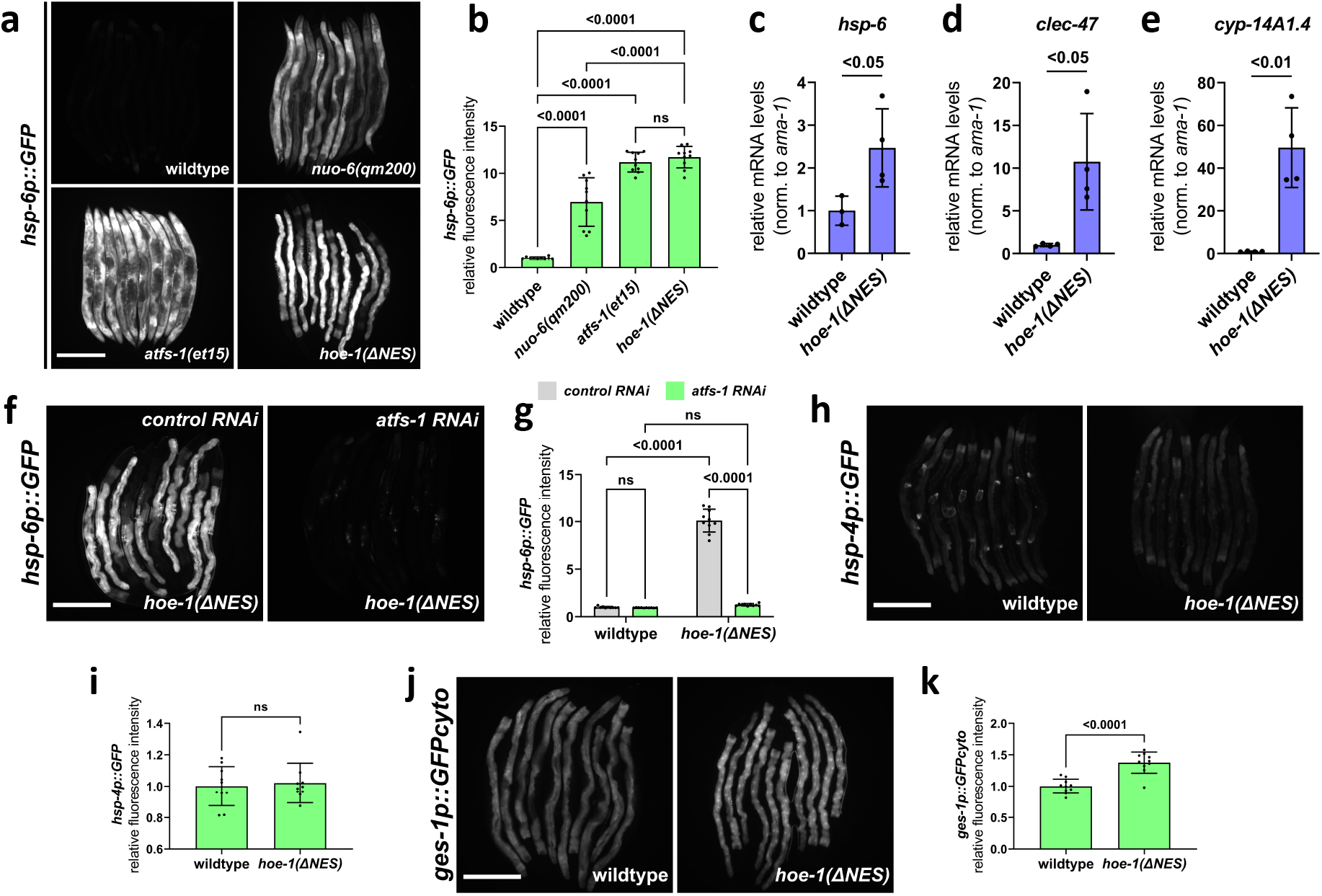
Nuclear export defective HOE-1 is sufficient to specifically activate UPR^mt^. **a**, Fluorescence images of UPR^mt^ reporter (*hsp-6p::GFP)* activation in day 2 adult wildtype, *nuo-6(qm200)*, *atfs-1(et15)*, and *hoe-1*(ΔNES) animals. Scale bar 200μm. **b**, Fluorescence intensity quantification of *hsp-6p::GFP* in individual day 2 adult wildtype, *nuo-6(qm200)*, *atfs-1(et15)*, and *hoe-1*(ΔNES) animals (n=10 for each condition, mean and SD shown, ordinary one-way ANOVA with Tukey’s multiple comparisons test). **c**, **d**, **e**, mRNA transcript quantification of *hsp-6*, *clec-47*, and *cyp-14A1.4*, respectively, in day 2 adult wildtype and *hoe-1*(ΔNES) animals normalized to RNA pol II, *ama-1*, mRNA levels (n=4 for each condition, mean and SD shown, two-tailed unpaired t-test). **f**, Fluorescence images of UPR^mt^ reporter (*hsp-6p::GFP)* activation in day 2 adult *hoe-1*(ΔNES) animals on control and *atfs-1 RNAi*. Scale bar 200μm. **g**, Fluorescence intensity quantification of *hsp-6p::GFP* in individual day 2 adult wildtype and *hoe-1*(ΔNES) animals on control and *atfs-1 RNAi* (n=10 for each condition, mean and SD shown, ordinary two-way ANOVA with Tukey’s multiple comparisons test). **h**, Fluorescence images of UPR^ER^ reporter (*hsp-4p::GFP*) activation in day 2 adult wildtype and *hoe-1*(ΔNES) animals. Scale bar 200μm. **i**, Fluorescence intensity quantification of *hsp-4p::GFP* in individual day 2 adult wildtype and *hoe-1*(ΔNES) animals (n=10 for each condition, mean and SD shown, two-tailed unpaired t-test). **j**, Fluorescence images of intestinal-specific basal protein reporter (*ges-1p::GFPcyto*) activation in day 2 adult wildtype and *hoe-1*(ΔNES) animals. Scale bar 200μm. **k**, Fluorescence intensity quantification of *ges-1p::GFPcyto* in individual day 2 adult wildtype and *hoe-1*(ΔNES) animals (n=10 for each condition, mean and SD shown, two-tailed unpaired t-test).

UPR^mt^ activation is characterized by the transcriptional upregulation of a suite of mitochondrial stress response genes that encode chaperone proteins, proteases, and detoxification enzymes that function to restore mitochondrial homeostasis ^14^. To interrogate the extent of UPR^mt^ induction in *hoe-1*(ΔNES) animals, we measured transcript levels of a diverse set of UPR^mt^ associated genes. We found that the UPR^mt^ genes encoding a chaperone protein (*hsp-6*), stress response involved C-type lectin (*clec-47*) and P450 enzyme (*cyp-14A4.1*) are all upregulated in *hoe-1*(ΔNES) animals **(Fig. 3c, d, e)**. These data support *hoe-1*(ΔNES) being sufficient to activate the UPR^mt^ transcriptional response.

UPR^mt^ activation is dependent upon the transcription factor ATFS-1 ^14,15^. Thus, we tested if UPR^mt^ reporter activation in *hoe-1*(ΔNES) animals is ATFS-1 dependent. Knockdown of *atfs-1* is sufficient to completely attenuate UPR^mt^ reporter activation in *hoe-1*(ΔNES) animals **(Fig. 3f, g)**, showing that UPR^mt^ induction by *hoe-1*(ΔNES) is ATFS-1 dependent.

### Compromising HOE-1 nuclear export specifically activates UPR^mt^

Changes in protein synthesis rates and associated protein folding capacity can broadly activate cellular stress response mechanisms ^43–45^. Given the role of *hoe-1* in tRNA maturation we questioned if the robust upregulation of UPR^mt^ in *hoe-1*(ΔNES) animals may be the result of compromised cellular proteostasis in general rather than specific activation of UPR^mt^. One stress response mechanism that is sensitive to global proteotoxic stress is the endoplasmic reticulum unfolded protein response (UPR^ER^) ^46^. We find that the UPR^ER^ reporter *hsp-4p::GFP* is not activated in *hoe-1*(ΔNES) animals **(Fig. 3h, i)**, suggesting that *hoe-1*(ΔNES) does not cause ER stress nor cellular proteotoxic stress. Additionally, a basal reporter of GFP that has been used to proxy general protein expression ^47^, *ges-1p::GFPcyto*, is only mildly upregulated in *hoe-1*(ΔNES) animals relative to wildtype **(Fig. 3j, k)**. Together these findings support that impaired nuclear export of HOE-1 specifically activates UPR^mt^.

### *hoe-1*(ΔNES) animals have elevated nuclear levels of UPR^mt^ transcription factors ATFS-1 and DVE-1

We tested if ATFS-1 accumulates in nuclei of *hoe-1*(ΔNES) animals by assessing the fluorescence intensity of ectopically-expressed mCherry-tagged ATFS-1 (*atfs-1p*::ATFS-1::mCherry) in wildtype and *hoe-1*(ΔNES) animals. *hoe-1*(ΔNES) animals have elevated nuclear accumulation of ATFS-1 relative to wildtype animals **(Fig. 4a and Ext. Data Fig. 5)**.

**Figure 4:**
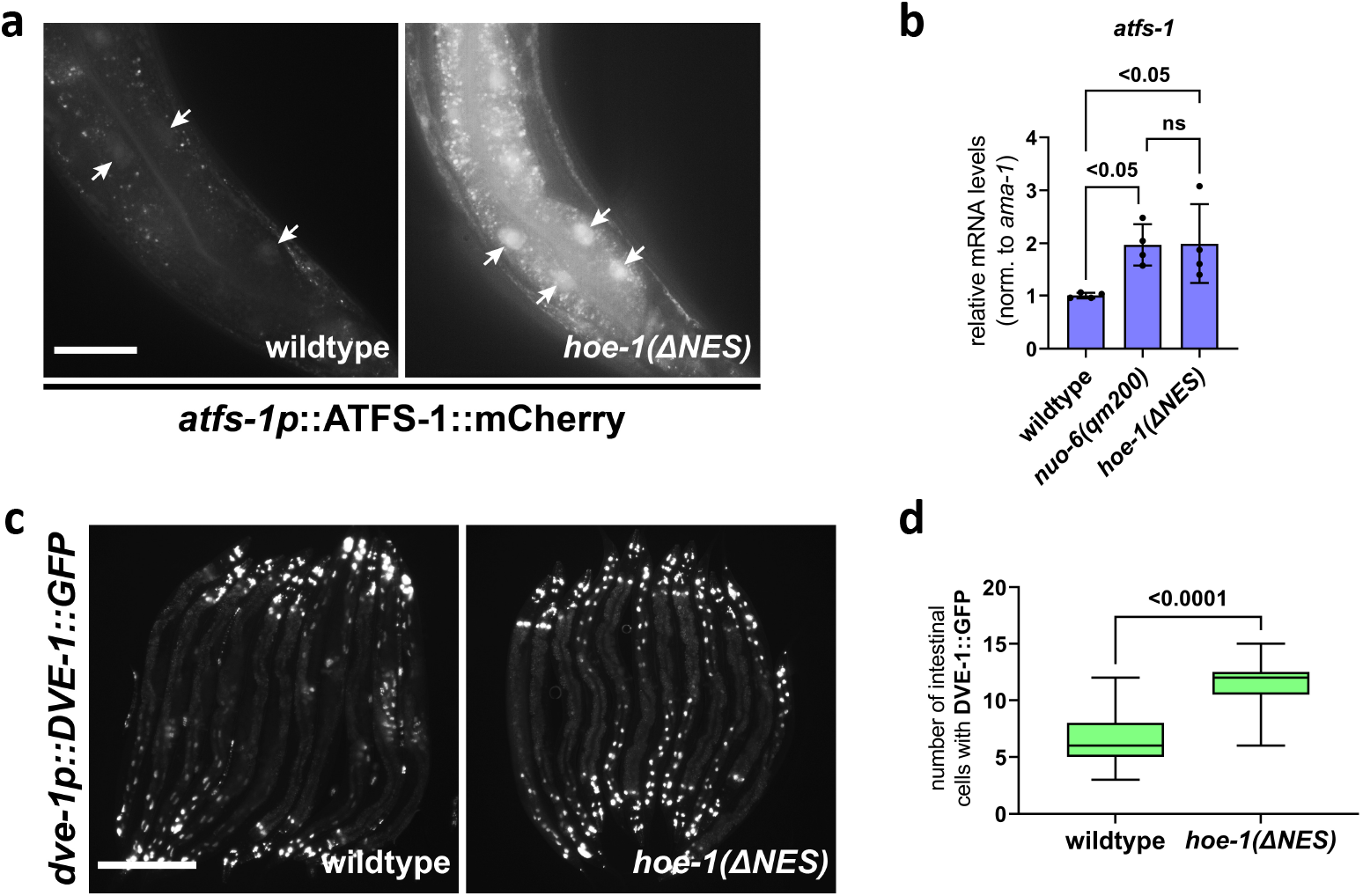
*hoe-1*(ΔNES) animals have increased nuclear accumulation of UPR^mt^ transcription factors ATFS-1 and DVE-1. **a**, Fluorescence images of *atfs-1::ATFS-1::mCherry* in the terminal intestine of day 2 adult wildtype and *hoe-1*(ΔNES) individuals (tip of the tail is in the bottom right corner of each panel). Arrows indicate intestinal nuclei. Scale bar 25μm. **b**, mRNA transcript quantification of *atfs-1* in day 2 adult wildtype, *nuo-6(qm200)*, and *hoe-1*(ΔNES) animals normalized to *ama-1* (n=4 for each condition, mean and SD shown, ordinary one-way ANOVA with Tukey’s multiple comparisons test). **c**, Fluorescence images of *dve-1p::DVE-1::GFP* in wildtype and *hoe-1*(ΔNES) day 2 adult animals. Scale bar 200μm. **d**, Number of intestinal cell nuclei with DVE-1::GFP puncta above brightness threshold of 25 in day 2 adult wildtype and *hoe-1*(ΔNES) animals (n=33 and 41 respectively, two-tailed unpaired t-test).

Given total ATFS-1 levels are elevated in *hoe-1*(ΔNES) animals, we questioned whether *atfs-1* is transcriptionally upregulated. We find that *atfs-1* mRNA levels are elevated in *hoe-1*(ΔNES) animals relative to wildtype but are comparable to that seen in *nuo-6(qm200)* animals **(Fig. 4b)** suggesting that transcriptional upregulation of ATFS-1 alone is insufficient to account for the robust activation of UPR^mt^ in *hoe-1*(ΔNES) animals.

The transcription factor DVE-1 is required for full UPR^mt^ activation ^48,49^. Thus we asked if DVE-1::GFP nuclear expression is higher in *hoe-1*(ΔNES) than in wildtype animals. We found that nuclear accumulation of DVE-1::GFP is higher in *hoe-1*(ΔNES) animals than wildtype **(Fig. 4c, d)**. Together these data suggest that UPR^mt^ induction in *hoe-1*(ΔNES) animals is a result of increased nuclear accumulation of UPR^mt^ transcription factors ATFS-1 and DVE-1.

### *hoe-1*(ΔNES) animals do not exhibit characteristics of mitochondrial stress

We questioned if UPR^mt^ activation in *hoe-1*(ΔNES) animals is a result of mitochondrial stress. Mitochondrial stress that triggers UPR^mt^ activation has widely been shown to delay development of *C. elegans* ^50,51^. Interestingly, *hoe-1*(ΔNES) animals develop similarly to wildtype animals rather than exhibiting slowed development like that seen in animals with mitochondrial stress (*nuo-6(qm200)*) **(Fig. 5a)**. Under conditions of mitochondrial stress, mitochondrial membrane potential (ΔΨm) is often compromised ^52^. Using TMRE staining to proxy ΔΨm indicates that there is no significant difference in ΔΨm between wildtype and *hoe-1*(ΔNES) L4 animals **(Fig. 5b)**. Taken together, these data suggest that *hoe-1*(ΔNES) induced UPR^mt^ is not a result of mitochondrial stress.

**Fig. 5:**
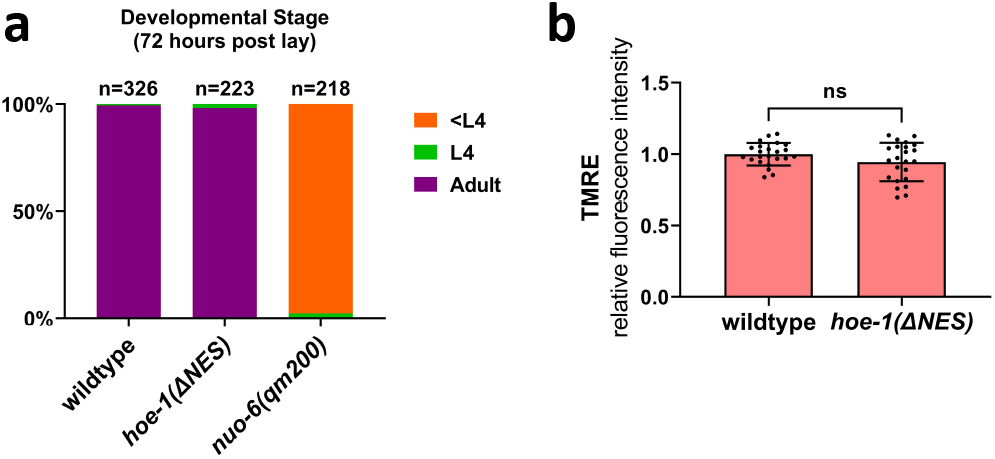
*hoe-1*(ΔNES) animals do not exhibit signs of mitochondrial stress. **a**, Developmental stage of wildtype, *nuo-6(qm200)*, and *hoe-1*(ΔNES) animals 72 hours post lay. **b**, Fluorescence intensity quantification of TMRE staining in individual L4 wildtype and *hoe-1*(ΔNES) animals (n=23 for each condition, mean and SD shown, two-tailed unpaired t-test).

### UPR^mt^ is activated by altered tRNA processing in *hoe-1*(ΔNES) animals

The canonical function of ELAC2 is to cleave 3’-trailer sequences from nascent tRNAs ^22–29^. The production of mature tRNAs begins with transcription of tRNA gene loci followed by sequential cleavage of 5’-leader and 3’-trailer sequences from immature tRNA transcripts. Following cleavage of 3’-trailer sequences, tRNAs can be transported to the cytosol by tRNA exportin ^53^.

Given that HOE-1 is restricted to the nucleus in *hoe-1*(ΔNES) animals, we reason that 3’-tRNA processing should be elevated. Thus, we questioned if UPR^mt^ induction in *hoe-1*(ΔNES) animals is a result of elevated 3’-tRNA processing. For the majority of tRNAs 5’-end processing by the RNase P complex is a prerequisite for 3’-end processing by ELAC2 ^54,55^. Thus, if increased 3’-tRNA end processing is responsible for UPR^mt^ activation, compromising 5’-end processing by RNAi against RNAse P should suppress *hoe-1*(ΔNES) induced UPR^mt^.

RNAi against a subunit of the RNase P complex, *popl-1*, attenuates UPR^mt^ induction in *hoe-1*(ΔNES) animals **(Fig. 6a, b)**. *popl-1* RNAi also attenuates both UPR^mt^ induced by *nuo-6(qm200)* **(Fig. 6c, d)** as well as basal induction of *ges-1p::GFPcyto* **(Fig. 6e, f)** but to a lesser extent than the attenuation seen in *hoe-1*(ΔNES) animals. These data suggest that *popl-1* RNAi may have a broad impact on protein expression but supports that elevated 3’-tRNA processing in *hoe-1*(ΔNES) animals is responsible for UPR^mt^ activation given that *popl-1* RNAi strongly attenuates *hoe-1*(ΔNES) induced UPR^mt^.

**Fig. 6:**
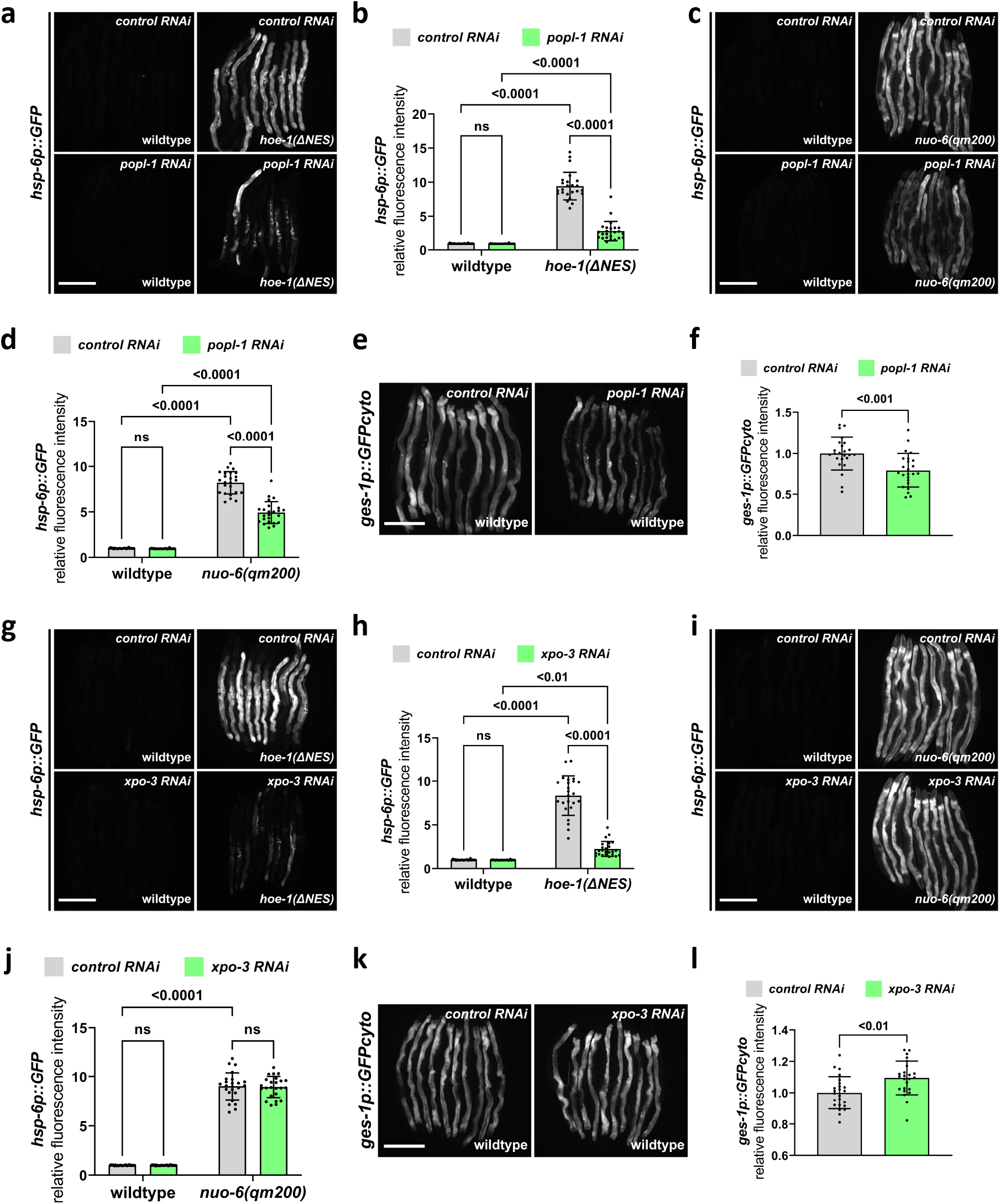
Nuclear export defective HOE-1 activates UPR^mt^ via altered tRNA processing. **a**, Fluorescence images of UPR^mt^ reporter (*hsp-6p::GFP)* activation in day 2 adult wildtype and *hoe-1*(ΔNES) animals on control and *popl-1 RNAi*. Scale bar 200μm. **b**, Fluorescence intensity quantification of *hsp-6p::GFP* in individual day 2 adult wildtype and *hoe-1*(ΔNES) animals on control and *popl-1 RNAi* (n=24 for each condition, mean and SD shown, ordinary two-way ANOVA with Tukey’s multiple comparisons test). **c**, Fluorescence images of UPR^mt^ reporter (*hsp-6p::GFP)* activation in day 2 adult wildtype and *nuo-6(qm200)* animals on control and *popl-1 RNAi*. Scale bar 200μm. **d**, Fluorescence intensity quantification of *hsp-6p::GFP* in individual day 2 adult wildtype and *nuo-6(qm200)* animals on control and *popl-1 RNAi* (n=24 for each condition, mean and SD shown, ordinary two-way ANOVA with Tukey’s multiple comparisons test). **e**, Fluorescence images of intestinal-specific basal protein reporter (*ges-1p::GFPcyto*) activation in day 2 adult wildtype animals on control and popl-1 RNAi. Scale bar 200μm. **f**, Fluorescence intensity quantification of *ges-1p::GFPcyto* in individual day 2 adult wildtype animals on control and *popl-1 RNAi* (n=24 for each condition, mean and SD shown, two-tailed unpaired t-test). **g**, Fluorescence images of UPR^mt^ reporter (*hsp-6p::GFP)* activation in day 2 adult wildtype and *hoe-1*(ΔNES) animals on control and *xpo-3 RNAi*. Scale bar 200μm. **h**, Fluorescence intensity quantification of *hsp-6p::GFP* in individual day 2 adult wildtype and *hoe-1*(ΔNES) animals on control and *xpo-3 RNAi* (n=24 for each condition, mean and SD shown, ordinary two-way ANOVA with Tukey’s multiple comparisons test). **i**, Fluorescence images of UPR^mt^ reporter (*hsp-6p::GFP)* activation in day 2 adult wildtype and *nuo-6(qm200)* animals on control and *xpo-3 RNAi*. Scale bar 200μm. **j**, Fluorescence intensity quantification of *hsp-6p::GFP* in individual day 2 adult wildtype and *nuo-6(qm200)* animals on control and *xpo-3 RNAi* (n=24 for each condition, mean and SD shown, ordinary two-way ANOVA with Tukey’s multiple comparisons test). **k**, Fluorescence images of intestinal-specific basal protein reporter (*ges-1p::GFPcyto*) activation in day 2 adult wildtype animals on control and *xpo-3 RNAi*. Scale bar 200μm. **l**, Fluorescence intensity quantification of *ges-1p::GFPcyto* in individual day 2 adult wildtype animals on control and *xpo-3 RNAi* (n=24 for each condition, mean and SD shown, two-tailed unpaired t-test).

Following 3’-end processing in the nuclei, tRNAs can be exported to the cytosol by tRNA exportin ^53^. To test if elevated levels of 3’-processed tRNAs are required in the cytosol to activate UPR^mt^ we asked if restricting tRNA nuclear export via RNAi against tRNA exportin, *xpo-3*, attenuates *hoe-1*(ΔNES) induced UPR^mt^. Strikingly, *xpo-3* RNAi robustly attenuates *hoe-1*(ΔNES) induced UPR^mt^ **(Fig. 6g, h)**. However, *xpo-3* RNAi does not attenuate *nuo-6(qm200)* induced UPR^mt^ **(Fig. 6i, j)** nor basal *ges-1p::GFP* levels **(Fig. 6k, l)** suggesting that *xpo-3* RNAi specifically suppresses *hoe-1*(ΔNES)-induced UPR^mt^. These data suggest that UPR^mt^ induction in *hoe-1*(ΔNES) animals is a result of increased 3’-tRNA processing and that these tRNA species are required in the cytosol to trigger UPR^mt^.

### *hoe-1*(ΔNES)-induced UPR^mt^ is not GCN2 or eIF2α dependent

Alteration to tRNA processing can activate cellular signaling pathways ^56^. One such pathway is the integrated stress response (ISR) in which uncharged tRNAs activate the kinase GCN2 which, in turn, phosphorylates the eukaryotic translation initiation factor, eIF2α. This inhibitory phosphorylation of eIF2α leads to upregulation of a select number of proteins including the transcription factor ATF4 ^57,58^. Interestingly, ATF4 and one of its targets, ATF5, are orthologs of ATFS-1 ^59^. Moreover, GCN2 and ISR in general have been shown to be responsive to mitochondrial stress ^3,10,11,60^. Thus, we questioned if UPR^mt^ activation by *hoe-1*(ΔNES) is mediated via GCN2 and eIF2α phosphorylation. We found that *hoe-1*(ΔNES)-induced UPR^mt^ is only slightly reduced in both a *gcn-2* null (*gcn-2(ok871)*) and an *eIF2α* non-phosphorylatable mutant (*eIF2α(S46A,S49A)*) background **(Fig. 7a, b)**. These data suggest that a mechanism independent of ISR must largely be responsible for UPR^mt^ activation by *hoe-1*(ΔNES).

**Fig. 7:**
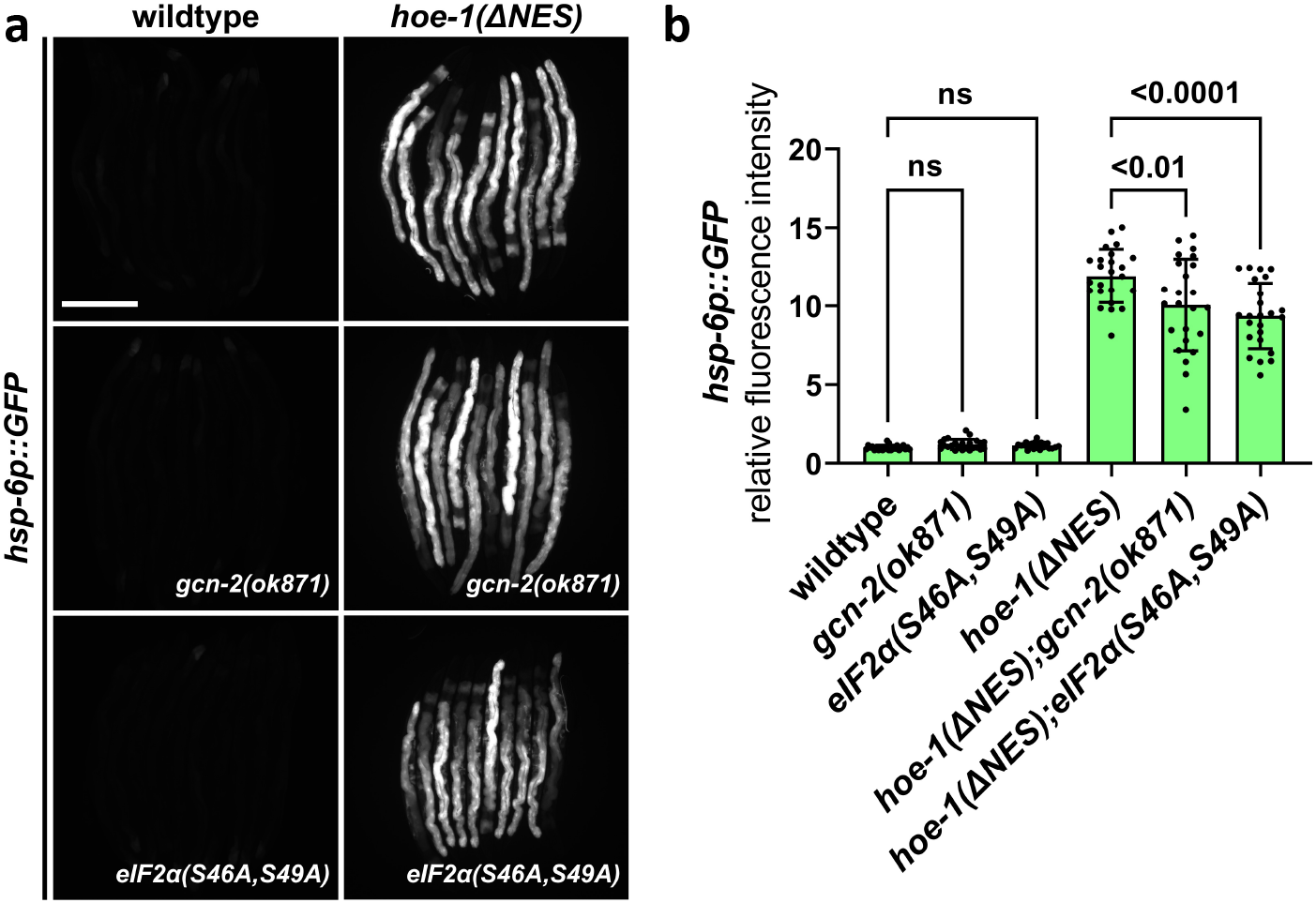
*hoe-1*(ΔNES)-induced UPR^mt^ is not *gcn-2* or *eIF2α* dependent. **a**, Fluorescence images of UPR^mt^ reporter (*hsp-6p::GFP)* activation in day 2 adult wildtype, *gcn-2(ok871)*, *eIF2α(S46A,S49A)*, *hoe-1*(ΔNES), *hoe-1*(ΔNES);*gcn-2(ok871)*, and *hoe-1*(ΔNES);*eIF2α(S46A,S49A)* animals. Scale bar 200μm. **b**, Fluorescence intensity quantification of *hsp-6p::GFP* in individual day 2 adult wildtype, *gcn-2(ok871)*, *eIF2α(S46A,S49A)*, *hoe-1*(ΔNES), *hoe-1*(ΔNES);*gcn-2(ok871),* and *hoe-1*(ΔNES);*eIF2α(S46A,S49A)* animals (n=24 for each condition, mean and SD shown, ordinary two-way ANOVA with Tukey’s multiple comparisons test).

### Nuclear HOE-1 is dynamically responsive to mitochondrial stress and negatively regulated by ATFS-1

To better understand the potential physiological implications of HOE-1 in UPR^mt^ we assessed the subcellular dynamics of HOE-1 during mitochondrial stress. We found that HOE-1::GFP nuclear levels are markedly diminished under mitochondrial stress induced by *nuo-6(qm200)* **(Fig. 8a, b)**. This observation runs contrary to the fact that compromising HOE-1 nuclear export is sufficient to induce UPR^mt^ **(Fig. 3a, b)**. A common feature of signaling pathways is negative regulation. Thus, we questioned if reduced nuclear HOE-1 is a result of negative feedback rather than a direct result of mitochondrial stress. Given that UPR^mt^ is induced when HOE-1 nuclear export is compromised, we assessed HOE-1::GFP status in a mitochondrial stress background when *atfs-1* is knocked down by RNAi. HOE-1 nuclear levels are significantly upregulated in nuclei of *nuo-6(qm200)* animals on *atfs-1* RNAi relative to *nuo-6(qm200)* animals on control RNAi, as well as both wildtype animals on control and *atfs-1* RNAi **(Fig. 8a, b)**. Moreover, total cellular HOE-1 levels are elevated under mitochondrial stress in an *atfs-1* RNAi background **(Fig. 8c, d).** These data suggest that HOE-1 is upregulated and accumulates in nuclei upon mitochondrial stress but is then negatively regulated by ATFS-1 once UPR^mt^ is activated.

**Fig. 8:**
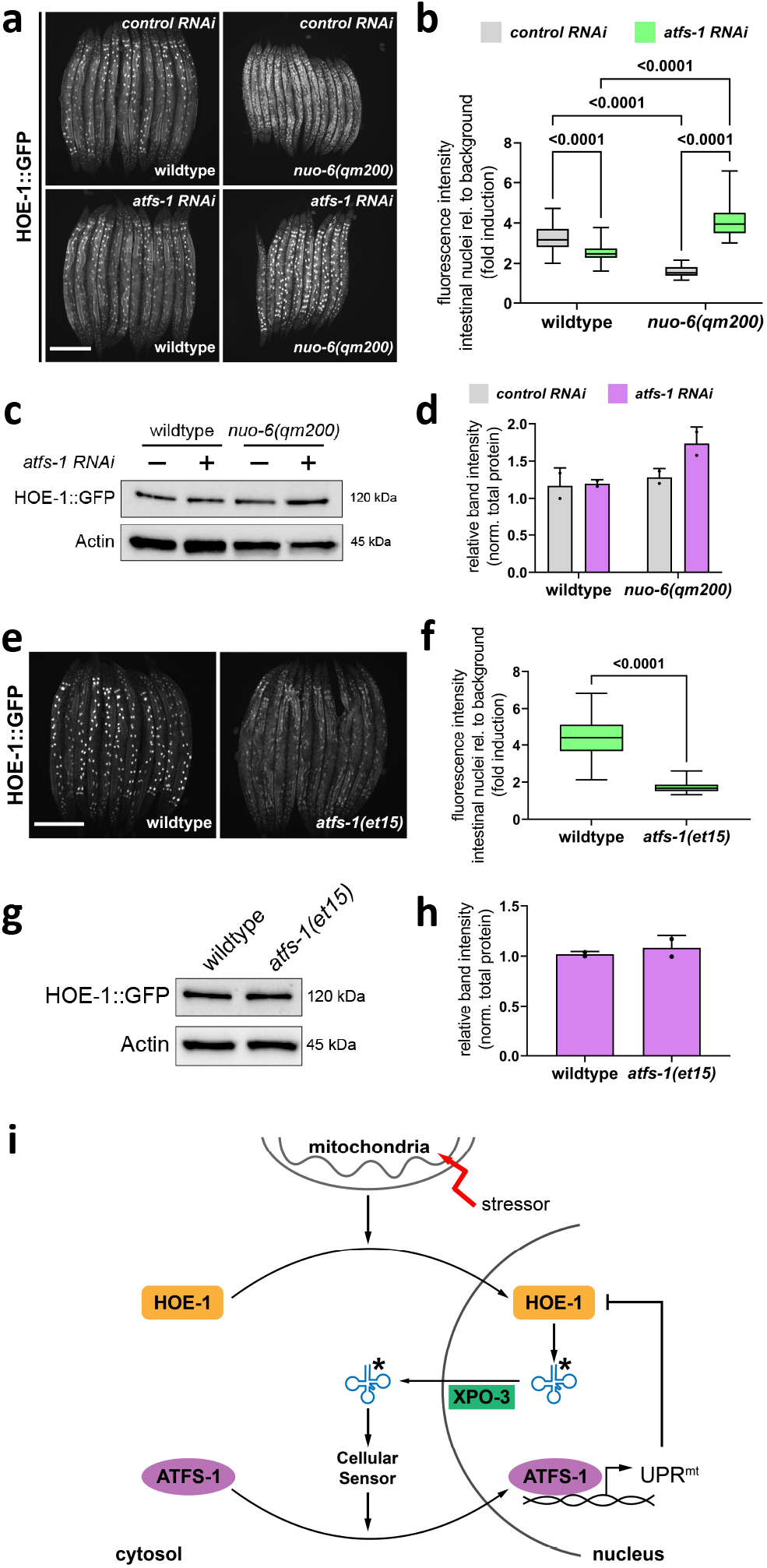
Nuclear HOE-1 levels are elevated during mitochondrial stress in the absence of ATFS-1 but decreased in the presence of ATFS-1. **a**, Fluorescence images of HOE-1::GFP in day 1 adult wildtype and *nuo-6(qm200)* animals on control and *atfs-1 RNAi.* Scale bar 200μm. **b**, Fluorescence intensity quantification of intestinal nuclei relative to background signal in day 1 adult wildtype and *nuo-6(qm200)* animals on control and *atfs-1 RNAi* (n=40 for each condition, ordinary two-way ANOVA with Tukey’s multiple comparisons test). **c**, Western blot for HOE-1::GFP and actin from day 1 adult wildtype and *nuo-6(qm200)* animals on control and *atfs-1 RNAi.* **d**, Quantification of HOE-1::GFP western blot band intensity from day 1 adult wildtype and *nuo-6(qm200)* animals on control and *atfs-1 RNAi* normalized to total protein (n=2 for each condition, mean and SD shown). **e**, Fluorescence images of HOE-1::GFP in day 1 adult wildtype and *atfs-1(et15)* animals. Scale bar 200μm. **f**, Fluorescence intensity quantification of intestinal nuclei relative to background signal in day 1 adult wildtype and *atfs-1(et15)* animals (n=40 for each condition, two-tailed unpaired t-test). **g**, Western blot for HOE-1::GFP and actin from day 1 adult wildtype and *atfs-1(et15)* animals. **h**, Quantification of HOE-1::GFP western blot band intensity from day 1 adult wildtype and *atfs-1(et15)* animals normalized to total protein (n=2 for each condition, mean and SD shown). **i**, Diagram of our model. Briefly, mitochondrial stress triggers the nuclear accumulation of HOE-1. Elevated nuclear HOE-1 results in altered tRNA processing (tRNAs*). tRNAs* are exported from the nucleus by XPO-3. In the cytosol tRNAs* trigger a signaling cascade that results in nuclear accumulation of ATFS-1. ATFS-1 drives UPR^mt^ activation which then negatively regulates nuclear HOE-1.

To further test if nuclear HOE-1 is negatively regulated by UPR^mt^ activation rather than by mitochondrial stress, we assayed HOE-1 localization in ATFS-1 gain-of-function animals (*atfs-1(et15)*). *atfs-1(et15)* constitutively activates UPR^mt^ in the absence of mitochondrial stress ^61^. Thus, we asked if *atfs-1(et15)* is sufficient to reduce nuclear HOE-1 levels. Indeed, nuclear HOE-1 levels are reduced in *atfs-1(et15)* animals relative to wildtype **(Fig. 8e, f)** while total HOE-1 protein levels are unperturbed **(Fig. 8g, h)**. These data further support that UPR^mt^ activation negatively regulates nuclear HOE-1.

## Discussion

Regulation of UPR^mt^ is not completely understood and elucidating this mechanism has broad implications for the understanding of mitochondrial dysfunction. Here we describe a novel mechanism by which mitochondrial stress is transduced to activate UPR^mt^ and how that response is regulated through a feedback mechanism **(Fig. 8i)**.

Multiple factors have been identified that are required for maximal activation of UPR^mt^. This includes the mitochondrial localized proteins, CLPP-1 protease and peptide transmembrane transporter HAF-1 ^15,48^. Additionally, the transcription factors ATFS-1 and DVE-1 along with the co-transcriptional activator UBL-5 are required for UPR^mt^ activation ^14–16,48,49,62^. Histone modifications, chromatin remodeling, and post-translational modifications of ATFS-1 are also involved in fully activating UPR^mt^ ^20,21,49,63^. We show for the first time that nuclear HOE-1 is required for maximal activation of UPR^mt^ as UPR^mt^ induction by various stressors is attenuated in a *hoe-1* RNAi, *hoe-1* null, and *hoe-1*(ΔNLS) backgrounds.

We show that loss of *hoe-1* results in varied attenuation of UPR^mt^ depending on how UPR^mt^ is activated. UPR^mt^ induction by RNAi (*cco-1* and *spg-7*) is robustly attenuated by loss of *hoe-1* while *nuo-6(qm200)*-induced UPR^mt^ is only modestly attenuated. RNAi by feeding works well in all tissues except neurons ^64,65^. Importantly, UPR^mt^ can be activated non-cell autonomously in the intestine by mitochondrial stress in neurons ^17–19^. Given that *hoe-1*(ΔNES)-induced UPR^mt^ is primarily restricted to the intestine, we reason that UPR^mt^ induced cell-autonomously in the intestine by RNAi is *hoe-1* dependent while neuron-to-intestine UPR^mt^ induction works primarily in a *hoe-1*-independent manner. These results further exemplify the complexity of UPR^mt^ signaling.

Mutations in ATFS-1 that compromise its mitochondrial localization have been shown to robustly induce UPR^mt^ in the absence of mitochondrial stress ^61^. We show that defective nuclear export of HOE-1 in *hoe-1*(ΔNES) adult animals is sufficient to activate UPR^mt^ seemingly independent of mitochondrial stress and just as robustly as the *atfs-1* gain-of-function mutation *atfs-1(et15)*. Furthermore, we show that ATFS-1 and DVE-1 nuclear levels are elevated in *hoe-1*(ΔNES) animals, thus likely facilitating the robust UPR^mt^ activation. Interestingly, we find that UPR^mt^ is not activated in *hoe-1*(ΔNES) animals until L4 larval stage. This could be a result of maternal deposition of wildtype HOE-1 from heterozygous mothers since the *hoe-1*(ΔNES) allele is balanced. Alternatively, this could be a biologically relevant timing phenomenon which requires further investigation.

HOE-1 functions in tRNA processing ^22–29^. Here we show that increased 3’-tRNA processing by HOE-1 is likely responsible for UPR^mt^ activation. Restricting HOE-1-dependent 3’tRNA trailer sequence cleavage indirectly by RNAi against RNAseP subunit *popl-1* attenuates *hoe-1*(ΔNES) induced UPR^mt^. Moreover, these tRNA species must be required in the cytosol to activate UPR^mt^ as RNAi against tRNA exportin *xpo-3* is sufficient to specifically attenuate *hoe-1*(ΔNES) induced UPR^mt^. Our findings herein are the first reported connection between altered tRNA processing and UPR^mt^ in *C. elegans*. There are outstanding questions regarding this relationship that we look forward to interrogating: 1) What is/are the tRNA species responsible for UPR^mt^ activation, and, 2) How does it signal UPR^mt^ induction? We reason that in *hoe-1*(ΔNES) animals there is increased 3’-end processing of tRNAs above wildtype levels given that HOE-1 is restricted to the nucleus, thus resulting in a greater pool of fully processed tRNAs. Following tRNA end processing, tRNAs undergo CCA addition at their 3’-ends. This modification is a prerequisite for aminoacylation of tRNAs to then be used for protein synthesis ^66^. Whether these steps are able to keep up with the presumed increased pool of 3’processed tRNAs we do not yet know. Thus, there are two broad types of tRNA species that could be involved in signaling: either an increase in fully matured, charged tRNAs, or accumulation of immature tRNA precursors. Indeed immature tRNA species are known to be used for signal transduction. In fact, uncharged tRNAs activate the eIF2α kinase, GCN2, resulting in the upregulation of ATFS-1 orthologs ATF4 and ATF5 ^57,58^. However, we show that *gcn-2* and eIF2α are not required for *hoe-1*(ΔNES)-induced UPR^mt^ activation suggesting that a different mechanism is responsible. It is possible that a tRNA-like RNA is responsible for UPR^mt^ induction in *hoe-1*(ΔNES) animals given that HOE-1 orthologs are capable of processing other structured RNAs ^29–32^. However, if this is the case, our data argue that such an RNA species would need to be transported to the cytosol by tRNA exportin. Non-tRNA transport by an ortholog of *xpo-3* has not been reported ^53^. Nonetheless, the question remains as to how such an RNA species induces UPR^mt^. Our data show that *hoe-1*(ΔNES) animals have elevated ATFS-1 and DVE-1 nuclear levels suggesting that the signaling mechanism downstream of HOE-1 must, at least in part, function to upregulate these transcription factors.

We show that nuclear HOE-1 is dynamically regulated by mitochondrial stress. In the presence of stress, nuclear HOE-1 levels are depleted. However, this is UPR^mt^ dependent as HOE-1 nuclear levels under mitochondrial stress are elevated above wildtype levels when UPR^mt^ is blocked by *atfs-1* RNAi. These data, paired with the fact that compromising HOE-1 nuclear export triggers UPR^mt^, lead us to hypothesize that upon mitochondrial stress, nuclear HOE-1 levels are elevated. This upregulation of nuclear HOE-1 elevates 3’-tRNA processing thereby triggering a signaling cascade that results in elevated nuclear ATFS-1 and DVE-1 and subsequent UPR^mt^ induction. Activated UPR^mt^ then negatively regulates HOE-1 nuclear levels thus providing a feedback mechanism to tightly control mitochondrial stress response. UPR^mt^ negative regulation of HOE-1 is further supported by our data showing that constitutive activation of UPR^mt^ by *atfs-1(et15)* is sufficient to reduce nuclear HOE-1 levels in the absence of mitochondrial stress. We look forward to identifying the players involved in attenuating nuclear HOE-1 levels upon UPR^mt^ induction.

In humans, mutations in the ortholog of HOE-1, ELAC2, are associated with both hypertrophic cardiomyopathy ^33–35^ and prostate cancer ^36–38^. Historically, it has been suggested that mutations in ELAC2 cause disease because of a loss of mature tRNA production. Our works suggests an intriguing alternative whereby ELAC2 mutations lead to altered tRNA processing that triggers aberrant stress response signaling resulting in disease state. Our system provides a convenient opportunity to interrogate these disease causing variants.

Taken together, our findings provide a novel mechanism—involving the tRNA processing enzyme HOE-1—by which mitochondrial stress is transduced to activate UPR^mt^ thus providing important insight into the regulation of mitochondrial stress response.

## Acknowledgements

We thank Lantana K Grub, Cassidy A Johnson, and Kristopher Burkewitz for their valuable feedback on the manuscript. Worm strain itSi001 was graciously shared with us by Sasha de Henau. Some strains were provided by the CGC, which is funded by NIH Office of Research Infrastructure Programs (P40 OD010440). This work was generously supported by R01 GM123260 (MRP) and by the support provided to JPH by the Training Program in Environmental Toxicology (T32ES007028). Confocal microscopy imaging was performed through the Vanderbilt Cell Imaging Shared Resource (supported by NIH grants CA68485, DK20593, DK58404, DK59637 and EY08126). Droplet Digital PCR to quantify transcript levels was performed through the Vanderbilt University Medical Center’s Immunogenomics, Microbial Genetics and Single Cell Technologies core.

## Author Contributions

Conceptualization, J.P.H. and M.R.P.; Methodology, J.P.H.; Validation, J.P.H.; Formal Analysis, J.P.H.; Investigation, J.P.H., C.V.P., and B.R.S.; Resources, M.R.P.; Writing – Original Draft, J.P.H; Writing – Review & Editing, J.P.H., C.V.P., and M.R.P.; Visualization, J.P.H.; Supervision, M.R.P.; Project Administration, J.P.H. and M.R.P.; Funding Acquisition, M.R.P.

## Methods

### Worm Maintenance

Worms were grown on nematode growth media (NGM) seeded with OP50 *E. coli* bacteria and maintained at 20°C.

### Mutants and Transgenic Lines

A complete list of *C. elegans* strains used can be found in supplemental table S1. All new mutant and transgenic strains generated via CRISPR/*Cas9* for this study were confirmed by Sanger sequencing.

### CRISPR/Cas9

CRISPR was conducted as previously described (Dokshin *et al. Genetics* 2018; Paix *et al. Genetics* 2015) using Alt-R® S.p. Cas9 Nuclease V3 (IDT #1081058) and tracrRNA (IDT #1072532). A complete list of crRNA and repair template sequences purchased from IDT can be found in supplemental table S2.

### Genetic Crosses

Strains resulting from genetic crosses were generated by crossing ~20 heterozygous males of a given strain to 5 – 8 L4 hermaphrodites of another strain (heterozygous males were generated by first crossing L4 hermaphrodites of that strain to N2 males). F1, L4 hermaphrodites were then cloned out and allowed to have self-progeny. F2 progeny were cloned out and once they had progeny were genotyped or screened (if fluorescent marker) for presence of alleles of interest. All genotyping primers were purchased from IDT and can be found in supplemental table S2.

### Fluorescence Microscopy

All imaging was done using Zeiss Axio Zoom V16 stereo zoom microscope unless otherwise indicated. Zeiss LSM510 META inverted confocal microscope was used for Fig. 2a and Ext. Data Fig. 1. Leica upright fluorescent compound microscope was used for Fig. 4a and Ext Data Fig. 5. For all imaging, worms were immobilized on 2% agar pads on microscope slides in ~2μl of 100mM levamisole (ThermoFisher #AC187870100) and then coverslip applied.

### Fluorescence Image Analysis

For whole animal fluorescence intensity quantification, total pixel and pixel intensity were quantified using imageJ and average pixel intensity for each worm was calculated. For DVE-1::GFP image analysis (Fig. 4d), brightness threshold was set to 25 in imageJ and then the number of gut cell nuclei that were saturated at this threshold were counted. For Fig. 7b and 7f, average pixel intensity was calculated within the bounds of gut cell nuclei and outside of the bounds of gut cell nuclei and then graphed as the ratio fluorescence intensity of nuclei to background.

### RNAi

RNAi by feeding was conducted as previously described (Gitschlag *et al. Cell Met.* 2016). Briefly, RNAi clones were grown overnight from single colony in 2 ml liquid culture of LB supplemented with 50 μg/ml ampicillin. To make 16 RNAi plates, 50 ml of LB supplemented with 50 μg/ml ampicillin was inoculated with 500 μl of overnight culture and then incubated while shaking at 37°C for 4 – 5 hours (to an OD550-600 of about 0.8). Cultures were then induced by adding 50 ml additional LB supplemented with 50 μg/ml ampicillin and 4mM IPTG and then continued incubating while shaking at 37°C for 4 hours. Following incubation, bacteria were pelleted by centrifugation at 3900 rpm for 6 minutes. Supernatant was decanted and pellets were gently resuspended in 4 ml of M9 buffer supplemented with 8mM IPTG. 250 μl of resuspension was seeded onto standard NGM plates containing 1mM IPTG. Plates were left to dry overnight and then used within 1 week. Bacterial RNAi feeder strains were all from Ahringer RNAi Feeding Library, grown from single colony and identity confirmed by Sanger sequencing. *atfs-1* (ZC376.7), *cco-1* (F26E4.9), *hoe-1* (E04A4.4), *popl-1* (C05D11.9), *spg-7* (Y47G6A.10), *xpo-3* (C49H3.10).

### Quantification of Gene Expression

cDNA was synthesized using Maxima H Minus First Strand cDNA Synthesis Kit, with dsDNase (ThermoFisher #K1682) according to manufacturer’s directions. Lysates for cDNA synthesis were made by transferring 10, day 2 adult worms to 10 μl of lysis buffer supplemented with 20mg/ml proteinase K and incubating at 65°C for 10 min, 85°C for 1 minute and 4°C for 2 minutes. Quantification of gene expression was performed using droplet digital PCR (ddPCR) with Bio-Rad QX200 ddPCR EvaGreen Supermix. Primers used for ddPCR can be found in supplemental table S2.

### Development Assay

50 gravid hermaphrodites were transferred to freshly seeded NGM plates and allowed to lay embryos for 2 hours. After 2 hours the adult hermaphrodites were removed. Plates with embryos were kept at 20°C. After 72 hours developmental stage of all progeny on the plate was scored.

### TMRE Staining

500 μl of 1mM TMRE (ThermoFisher #T669) was supplemented on top of fresh seeded standard NGM plates and allowed to dry overnight. The following day young L4 animals were transferred to TMRE plates. After 5 hours animals were transferred from TMRE plates to seeded standard NGM plates for 1 hour. Animals were then individually imaged in the red channel for fluorescence intensity. Whole animal fluorescence intensity was quantified as described above.

### Western Blot

50 adult worms were transferred into a tube containing 20 μl of M9 Buffer. Then, 20 μl of 2x Laemmli Buffer (BioRad #161-0737) supplemented with 2-mercaptoethanol (i.e. βME) was added to worm suspension and gently pipette up and down 5 times to mix. Worms were lysed at 95°C for 10min in thermocycler followed by ramp down to room temperature (25°C). Lysates were then pipetted up and down 10 times to complete disrupt and homogenize suspension. 20 μl of lysate was loaded onto precast Mini-PROTEAN TGX Stain-Free Gel (BioRad #4568045). Gel was run for 30 min at 100V and then an additional 40 – 45 min at 130V in 1x Tris/Glycine/SDS Running Buffer (BioRad #1610732). Following electrophoresis gel was activated and imaged for total protein. Gel was equilibrated in Trans-Blot® Turbo™ Transfer Buffer (BioRad #10026938) and transferred to activated and equilibrated Trans-Blot® Turbo™ LF PVDF Membrane (BioRad #10026934) for 7 min at 2.5A/25V on Trans-Blot® Turbo™ Transfer System. Following transfer, stain-free membrane was imaged for total protein. Membrane was then blocked in 5% milk in TBST for 2 hour rocking at room temperature. Following blocking, membrane was incubated in primary antibody overnight rocking at 4°C. Mouse monoclonal anti-β-actin (Santa Cruz Biotechnology #sc-47778) or mouse monoclonal anti-GFP (#sc-9996) were used at a dilution of 1:2500 in 5% milk in TBST. The following day the membrane was washed 3 times for 5 min each with TBST and then incubated with HRP-conjugated goat anti-mouse antibody (sc-2005) at 1:2000 in 5% milk in TBST for 2 hours at room temperature. Membrane was again washed 3 times for 5 min each with TBST. Membranes were then incubated for 5 minutes in Clarity™ Western ECL Substrate (BioRad #1705060) and immediately imaged on a BioRad ChemiDoc™ MP imager. Band intensity was quantified using imageJ.

**Extended Data Fig. 1:**
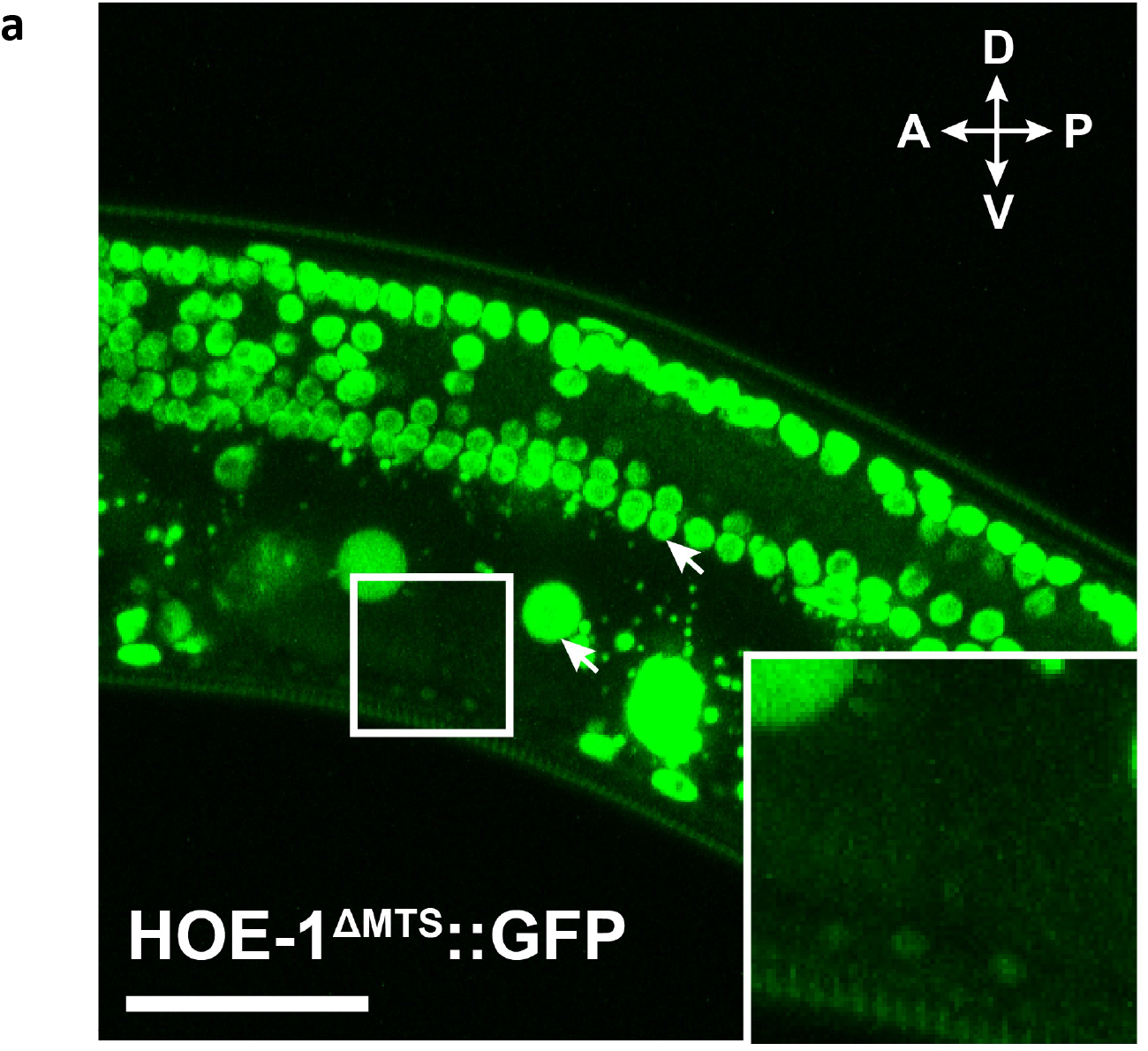
*hoe-1*(ΔMTS) allele ablates HOE-1 mitochondrial localization. **a**, Partial z-stack maximum intensity projection fluorescence image of *hoe-1*(ΔMTS) tagged with GFP (HOE-1^ΔMTS^::GFP) in adult germline. Orientation compass: anterior (A), posterior (P), dorsal (D), ventral (V). Arrows indicate nuclei. Area in the white square magnified in inset shows absence of mitochondrial signal. Scale bar 25μm.

**Extended Data Fig. 2:**
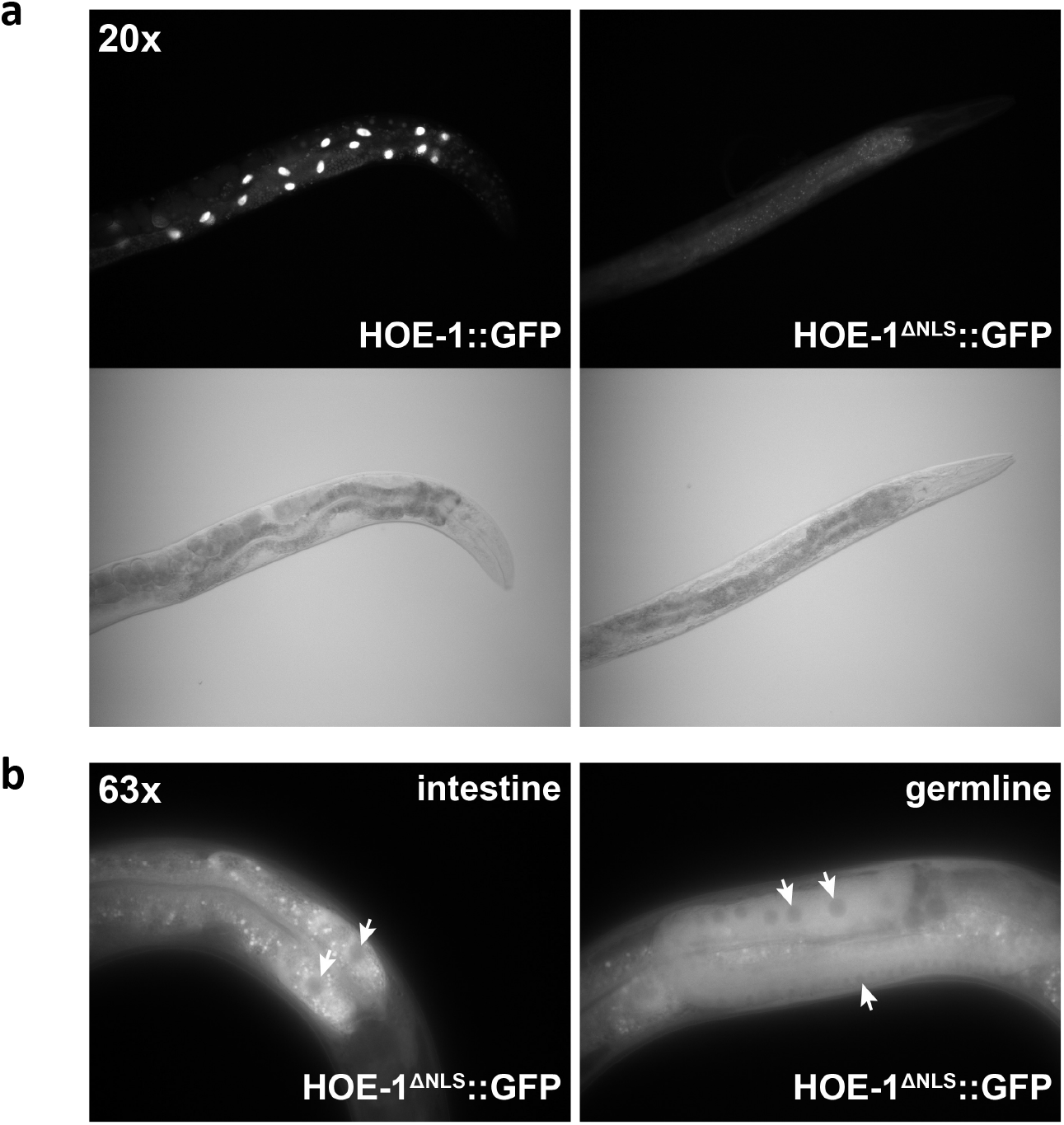
*hoe-1*(ΔNLS) allele ablates nuclear HOE-1 localization. **a**, Fluorescence and bright-field images of individual adults expressing HOE-1::GFP and HOE-1::GFP in a *hoe-1*(ΔNLS) background (HOE-1^ΔNLS^::GFP) under 20x objective on compound microscope. Large fluorescent puncta are intestinal cell nuclei. **b**, Fluorescence images of HOE-1^ΔNLS^::GFP intestine and germline under 63x objective on compound microscope. Arrows indicate nuclei devoid of GFP signal.

**Extended Data Fig. 3:**
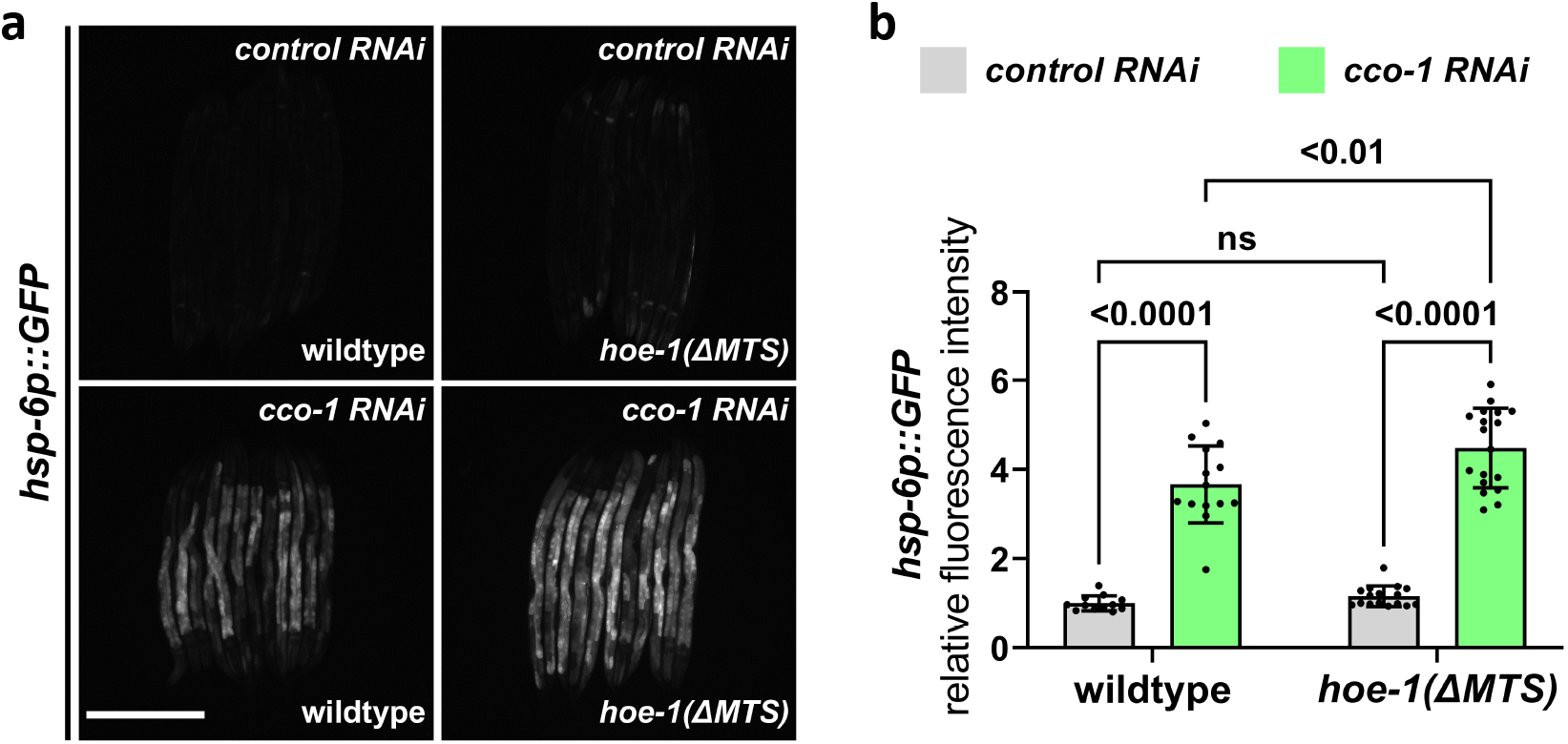
*hoe-1*(ΔMTS) does not attenuate *cco-1 RNAi-*induced UPR^mt^. **a**, Fluorescence images of UPR^mt^ reporter (*hsp-6p::GFP*) in L4 wildtype and *hoe-*1(ΔMTS) animals on control and *cco-1 RNAi*. Scale bar 200μm. **b**, Fluorescence intensity quantification of *hsp-6p::GFP* in individual L4 wildtype and *hoe-1*(ΔMTS) animals on control and *cco-1 RNAi* (n=12,14,16, and 18 respectively, mean and SD shown, ordinary two-way ANOVA with Tukey’s multiple comparisons test).

**Extended Data Fig. 4:**
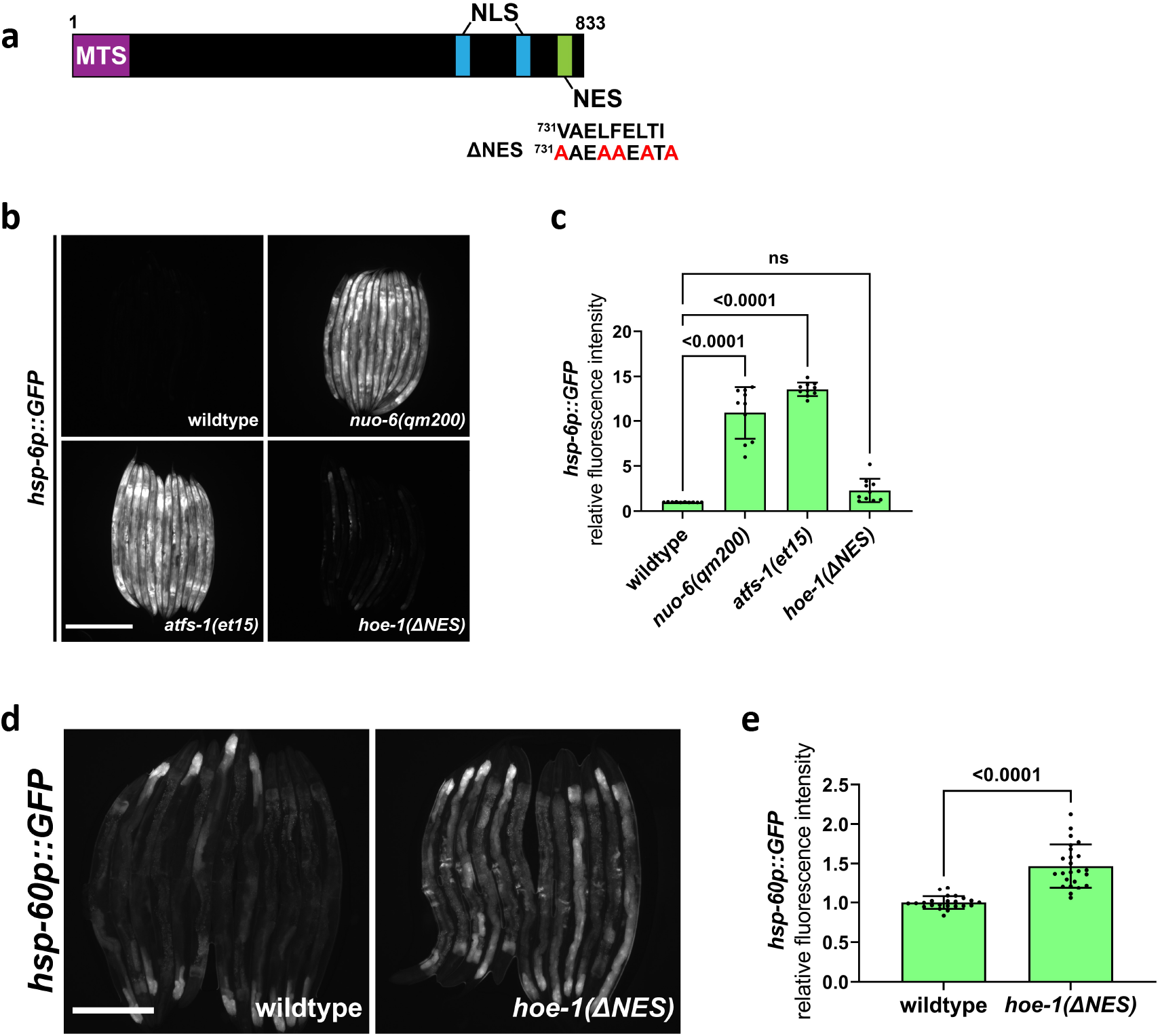
Nuclear export defective HOE-1 is sufficient to activate UPR^mt^. **a**, Schematic of HOE-1 protein showing the mitochondrial targeting sequence (MTS), nuclear localization signals (NLS) and nuclear export signal (NES). *hoe-1*(ΔNES) mutant generated by changing the strong hydrophobic residues of NES to alanines (^731^VAELFELTI^739^>^731^AAEAAEATA^739^) **b**, Fluorescence images of UPR^mt^ reporter (*hsp-6p::GFP)* activation in L4 wildtype, *nuo-6(qm200)*, *atfs-1(et15)*, and *hoe-1*(ΔNES) animals. Scale bar 200μm. **c**, Fluorescence intensity quantification of *hsp-6p::GFP* in individual L4 wildtype, *nuo-6(qm200)*, *atfs-1(et15)*, and *hoe-1*(ΔNES) animals (n=10 for each condition, mean and SD shown, ordinary one-way ANOVA with Tukey’s multiple comparisons test). **d**, Fluorescence images of UPR^mt^ reporter (*hsp-60p::GFP)* activation in day 2 adult wildtype and *hoe-1*(ΔNES) animals. Scale bar 200μm. **e**, Fluorescence intensity quantification of *hsp-60p::GFP* in individual day 2 adult wildtype and *hoe-1*(ΔNES) animals (n=24 for each condition, mean and SD shown, two-tailed unpaired t-test).

**Extended Data Fig. 5:**
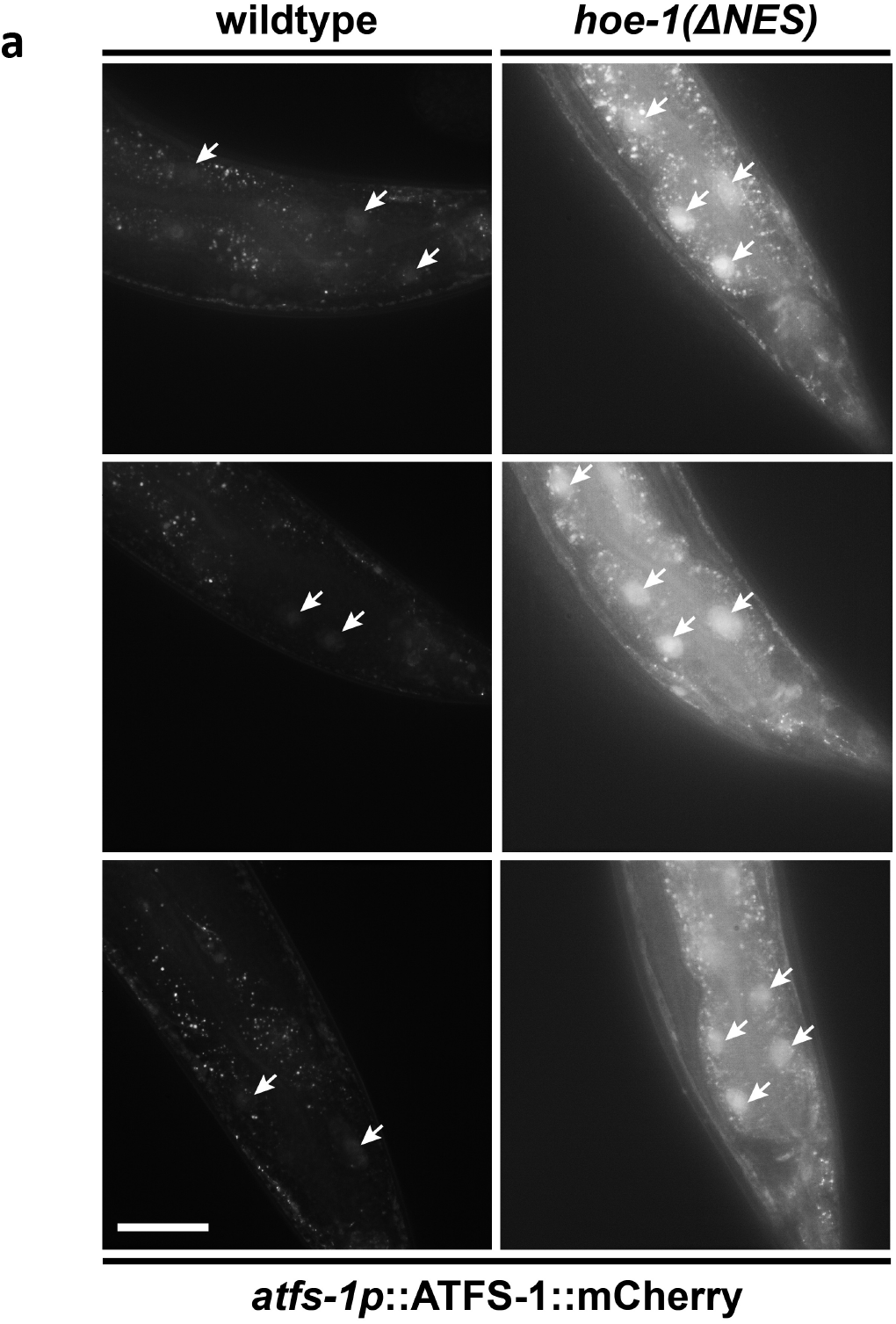
*hoe-1*(ΔNES) animals have increased nuclear ATFS-1 accumulation. **a**, Fluorescence images of *atfs-1::ATFS-1::mCherry* in the terminal intestine of day 2 adult wildtype and *hoe-1*(ΔNES) individuals (tip of the tail is in the right/bottom right corner of each panel). Arrows indicate intestinal nuclei. Scale bar 25μm.

**Extended Data Fig. 6:**
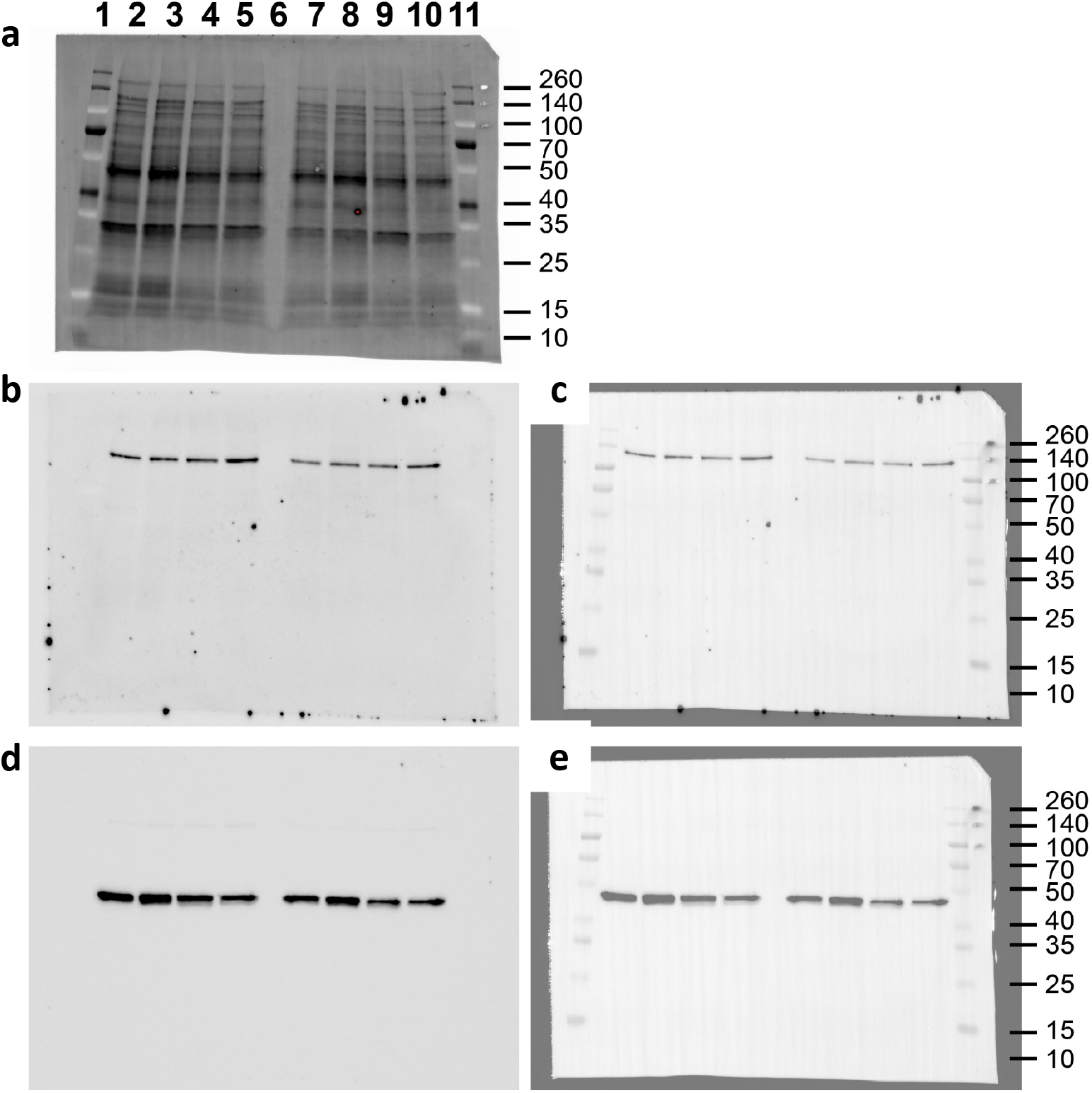
Blots for wildtype and *nuo-6(qm200)* animals on control and *atfs-1 RNAi* (Fig. 5c&d). All panels are the same membrane. **a**, Image of stain-free blot for total protein from day 1 adult wildtype and *nuo-6(qm200)* animals on control and *atfs-1 RNAi.* Two biological replicates of each condition: lane # 2&7 wildtype on control RNAi, 3&8 wildtype on *atfs-1 RNAi*, 4&9 *nuo-6(qm200)* on control RNAi, and 5&10 *nuo-6(qm200)* on *atfs-1 RNAi*. Lane # 1&11 BR Spectra Protein Ladder – ladder bands in kDa denoted. Lane # 5 empty. **b**, Chemiluminescence image of blot for HOE-1::GFP using GFP primary antibody. **c**, Composite image of chemiluminescence and colorimetric images of blot for HOE-1::GFP to show bands relative to ladder. **d**, Chemiluminescence image of blot for actin using β-actin primary antibody. **e**, Composite image of chemiluminescence and colorimetric images of blot for actin to show bands relative to ladder.

**Extended Data Fig. 7:**
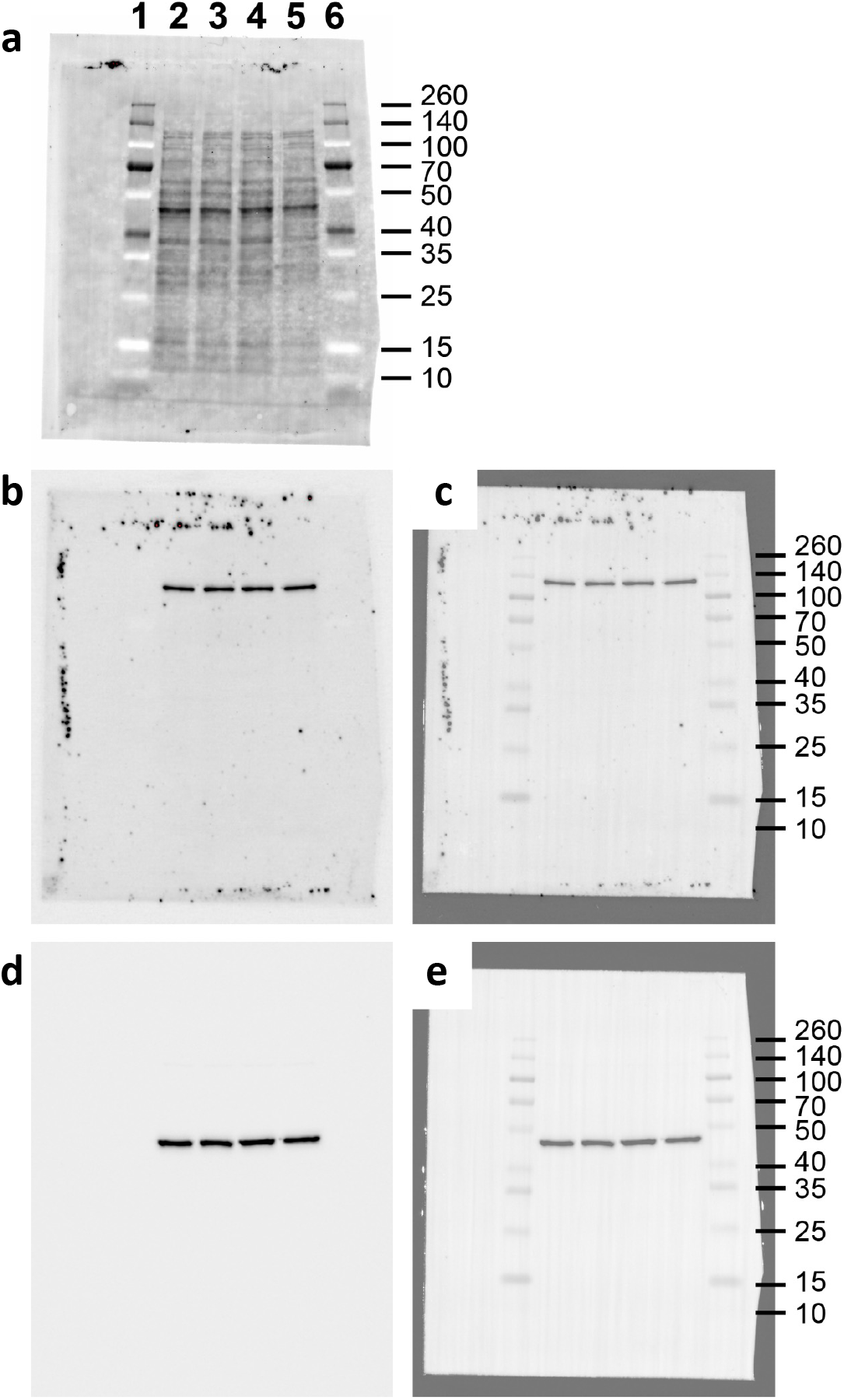
Blots for wildtype and *atfs-1(et15)* animals (Fig. 5g&h). All panels are the same membrane. **a**, Image of stain-free blot for total protein from day 1 adult wildtype and *atfs-1(et15)* animals. Two biological replicates of each condition: lane #2&4 wildtype and #3&5 *atfs-1(et15)*. Lane # 1&6 BR Spectra Protein Ladder – ladder bands in kDa denoted. **b**, Chemiluminescence image of blot for HOE-1::GFP using GFP primary antibody. **c**, Composite image of chemiluminescence and colorimetric images of blot for HOE-1::GFP to show bands relative to ladder. **d**, Chemiluminescence image of blot for actin using β-actin primary antibody. **e**, Composite image of chemiluminescence and colorimetric images of blot for actin to show bands relative to ladder.

