## Supplementary Tables for "A tRNA processing enzyme is a central regulator of the mitochondrial unfolded protein response"

**Supplemental S1**

| ***C. elegans* strains** | **Source** | **Strain Name** |
|---|---|---|
| *C. elegans* wild isolate | Caenorhabditis Genetics Center | N2 |
| *nuo-6(qm200)I* | Caenorhabditis Genetics Center | MQ1333 |
| *tmC25[unc-5(tmIs1241)] IV* | Caenorhabditis Genetics Center | FX30203 |
| *zcIs4 [hsp-4::GFP] V* | Caenorhabditis Genetics Center | SJ4005 |
| *zcIs9 [hsp-60::GFP + lin-15(+)]V* | Caenorhabditis Genetics Center | SJ4058 |
| *zcIs13[hsp-6p::GFP] V* | Caenorhabditis Genetics Center | SJ4100 |
| *zcIs18[ges-1::GFP(cyt)]* | Caenorhabditis Genetics Center | SJ4144 |
| *zcIs39[dve-1p::dve-1::GFP]II* | Caenorhabditis Genetics Center | SJ4197 |
| *atfs-1(et15)V, zcIs13[hsp-6p::GFP]V* | This study | MRP228 |
| *nuo-6(qm200)I; zcIs13[hsp-6p::GFP]V* | This study | MRP232 |
| *hoe-1(mpt13[M1A])IV; zcIs13(hsp-6p::GFP)V* | This study | MRP257 |
| *hoe-1(mpt14[HOE-1::GFP])IV* | This study | MRP258 |
| *hoe-1(mpt20[hoe-1[M1A]::GFP])IV* | This study | MRP264 |
| *hoe-1(mpt31[null])/tmC25 IV; zcIs13[hsp-6p::GFP]V* | This study | MRP314 |
| *hoe-1(mpt14[HOE-1::GFP])IV; atfs-1(et15)V* | This study | MRP330 |
| *nuo-6(qm200) I; hoe-1(mpt31[null])/tmC25 IV; zcIs13[hsp-6p::GFP]V* | This study | MRP333 |
| *hoe-1(mpt51[∆NLS])/tmC25 IV; zcIs13[hsp-6p::GFP]V* | This study | MRP335 |
| *hoe-1(mpt61[ΔNLS::GFP])/tmC25 IV* | This study | MRP341 |
| *nuo-6(qm200)I; hoe-1(mpt14[HOE-1::GFP])IV* | This study | MRP345 |
| *hoe-1(mpt67[ΔNES])/tmC25 IV* | This study | MRP372 |
| *hoe-1(mpt67[ΔNES])/tmC25 IV; zcIs13[hsp-6p::GFP]V* | This study | MRP379 |
| *hoe-1(mpt67[ΔNES])/tmC25 IV; zcIs18[ges-1::GFP(cyt)]* | This study | MRP384 |
| *hoe-1(mpt67[ΔNES])/tmC25 IV; zcIs4 [hsp-4::GFP] V* | This study | MRP393 |
| *ltSi001 II; hoe-1(mpt67[ΔNES])/tmC25 IV* | This study | MRP396 |
| *gcn-2(ok871) II; zcIs13[hsp-6p::GFP]V* | This study | MRP398 |
| *gcn-2(ok871) II; hoe-1(mpt67[ΔNES])/tmC25 IV; zcIs13[hsp-6p::GFP]V* | This study | MRP411 |
| *nuo-6(qm200) I; hoe-1(mpt51[∆NLS])/tmC25 IV; zcIs13[hsp-6p::GFP]V* | This study | MRP413 |
| *zcIs39[dve-1p::dve-1::GFP]II; hoe-1(mpt67[ΔNES])/tmC25 IV* | This study | MRP461 |
| *eIF-2alpha(mpt97) I; hoe-1(mpt67)/tmC25 IV; zcIs13[hsp-6p::GFP]V* | This study | MRP500 |
| *eIF-2alpha(mpt97[S46A,S49A]) I; zcIs13[hsp-6p::GFP]V* | This study | MRP503 |
| *hoe-1(mpt67[ΔNES])/tmC25 IV; zcIs9 [hsp-60::GFP + lin-15(+)]V* | This study | MRP515 |
| *ltSi001 [ttTi5605, Patfs-1::atfs-1::mCherry::atfs-1_3'UTR, cb-unc119(+)]II* | Sasha de Henau | |

**Supplemental S2. Oligonucleotides**

| | use/target | sequence |
|---|---|---|
| ***crRNAs*** | | |
| | dpy-10 (Arribere et al. 2014, Genetics) | GCTACCATAGGCACCACGAG |
| crRNA_JH07 | hoe-1(ΔMTS) / 5' for hoe-1(null) | CGAGGAAAACTGTAGAGAAT |
| crRNA_JH09 | hoe-1::GFP / 3' for hoe-1(null) | TGAAAAGCGAGGTCAATTAA |
| crRNA_JH18 | hoe-1(ΔNLS) | ACACCTCCCGGGAGCCCTGG |
| crRNA_JH25 | hoe-1(ΔNES) | TTACAGAGAAGTATTCGTGG |
| crRNA_JH29 | eIF2alpha(S46A,S49A) | TCACTGAACGGATACGACGA |
| ***ssDNA repairs*** | | |
| | dpy-10 (Arribere et al. 2014, Genetics) | CACTTGAACTTCAATACGGCAAGATGAGAATGACTGGAAACCGTACCGCATGCGGTGCCTATGGTAGCGGAGCTTCACATGGCTTCAGACCAACAGCCTAT |
| repair_JH007 | hoe-1(ΔMTS) | aattcaaataaatttcagctgaaagctcgaagctgGCTCTCGGAGCGATTGCGAGGAAAACTGTCGAAAATCGGATTCTgtaagggtttgtgagtggtccccc |
| repair_JH011 | hoe-1(null) | tgATGCTCGGAGCGATTGCGAGGAAAACTGTAGAGTAAAGGCTTGAgtgactggccacattcaacattaa |
| repair_JH016 | hoe-1(ΔNLS) | GTTGATATCTCCAGGTATCCACTAACACCTCCCGGCAGTCCGGGAGGGCCTCCTGGAGCTGCTCCGGCTCTTCCGAGCCCACATCTCCCGCCGAGCAGAGATGT |
| repair_JH025 | hoe-1(ΔNES) | aaataaataaatttataaatttacagAGAAGTATTTGCCGCTGAAGCGGCTGAAGCGACCGCTAAAAAAGAACAACGGGTTTTGAAAGACAAGGAATT |
| repair_JH032 | eIF2alpha(S46A,S49A) | aattaaatcatttttcagAGGGTATGATCCTGCTCGCTGAGCTCGCTCGACGTCGTATCCGTTCAGTGAACAAGCTAATCCGTGTCG |
| ***primers*** | | |
| **for gene expression** | | |
| JH037_F | hsp-6 cDNA | CGCTGGAGATAAGATCATCGCTG |
| JH038_R | | CGTTGGTGGACTTGACCTCG |
| JH039_F | ama-1 cDNA | GTACAATGCGGATTTCGATGGAGATG |
| JH040_R | | GGAGTGATGAGTTGTCTCGGC |
| JH111_F | atfs-1cDNA | CGAATAGATGAGTACGGAATGCCAC |
| JH112_R | | GTGATTCCATGCCTCCCAATCG |
| JH267_F | clec-47 cDNA | CCTGTGCTTGGGCAGATTCG |
| JH268_R | | CCCTGCTGTCTCAACAAATACACAG |
| JH269_F | cyp-14A1 cDNA | GGGTCTCATCAGTGCAAAGAGG |

| | | |
|---|---|---|
| JH270_R | | GGATCTTCAAACACTGTGTCATTTCTCATC |

**for genotyping**

| | | |
|---|---|---|
| JH133_F | nuo-6(qm200) | CCGAATTTCTGCCAGGACATGAATAC |
| JH134_R | follow w/ digest: TaqI cuts mutant | CCCTTGAACTCTCTCGAAAGTCAG |
| JH183_F | hoe-1(ΔMTS) | GGCATCTCCATCTCTCCCC |
| JH184_R | follow w/ digest: Cac8I cuts mutant | CCCGTTTTTTCTCAAAACCTGGAG |
| JH193_wt_F | hoe-1::GFP | GGTGTGGCGATGGATATGTTGAG |
| JH194_mt_F | | GGAAGCGTTCAACTAGCAGACC |
| JH195_R | | GCCTGTATATTTGGTTTTACCGCTAAATACC |
| JH218_mt_F | hoe-1(null) | GCTGAAAGCTCGAAGCTGATGC |
| JH219_wt_F | | GCCATAATGACCGAATATAACTGTGCTCG |
| JH220_R | | GCTAAATACCTAACGAGACCCAACGG |
| JH237_F | hoe-1(ΔNLS) | GCTGTCGGATAAAGCGAAGGAATTG |
| JH238_R | follow w/ digest: BsmA1 cuts WT | GGGGTTTTGGTGTCATATCTACCTCATG |
| JH295_mt_F | gcn-2(ok871) | GCTGATAGTGTATTTGAGAGATCCACTGC |
| JH296_wt_F | | CGGGCATCCACATGAGAGATGG |
| JH297_R | | GTTGTTCGCAATCGTGCTCGG |
| JH300_F | hoe-1(ΔNES) | CCCATTTTTAGGTAGATTTCAGATTCAGTATTCAGG |
| JH301_R | follow w/ digest: TaqI cuts WT | CGCTAAATACCTAACGAGACCCAACG |
| JH311_F | eIF2alpha(S46A,S49A) | GGCACGGAAAAAAGGGAAAAATTCG |
| JH312_R | follow w/ digest: AatII cuts mutant | GCACTAATATCACACTCATCCAAGATC |
| JH321_F | itSi001 (mCherry specific primers) | GGCATGAGTTTGAAATTGAAGGTGAAGG |
| JH322_R | | CATTGTACGCTCCTGGCAGC |
| MP397_F | atfs-1(et15) | GAAACCGCCTCCTTTCGCCTTTTG |
| MP398_R | follow w/ digest: AatII cuts WT | GACTTCATCGTCGTCCATGGGTACG |

**for dsDNA repair**

| | | |
|---|---|---|
| JH199_F | primers to amplify GFP from pJA245 | GAAAGACAAGGAATTGTCTGAAAAGCGAGGTCAATTGAAAGCTGGAGGTGGAGGTGGAGCTATGA GTAAAGGAGAAGAACT |
| JH200_R | include HA for hoe-1 and gly5A linker | AAAGTGAATTAATGTTGAATGTGGCCAGTCACTCATTATTTGTATAGTTCGTCCATGC |
